# HDAC8-mediated inhibition of EP300 drives a neural crest-like transcriptional state that increases melanoma brain metastasis

**DOI:** 10.1101/2022.10.12.511971

**Authors:** Michael F. Emmons, Richard L. Bennett, Alberto Riva, Chao Zhang, Robert Macaulay, Daphne Dupéré-Richér, Bin Fang, Edward Seto, John M. Koomen, Jiannong Li, Y. Ann Chen, Peter A. Forsyth, Jonathan D. Licht, Keiran S.M. Smalley

**Affiliations:** Department of Tumor Biology, Moffitt Cancer Center, 12902 Magnolia Drive, Tampa, Florida; Department of Neuro-Oncology, Moffitt Cancer Center, 12902 Magnolia Drive, Tampa, Florida; Proteomics & Metabolomics Core, Moffitt Cancer Center, 12902 Magnolia Drive, Tampa, Florida; Department of Molecular Oncology, Moffitt Cancer Center, 12902 Magnolia Drive, Tampa, Florida; Department of Bioinformatics and Biostatistics, Moffitt Cancer Center, 12902 Magnolia Drive, Tampa, Florida; Department of Cutaneous Oncology, Moffitt Cancer Center, 12902 Magnolia Drive, Tampa, Florida; University of Florida Health Sciences Center, Gainesville, Florida; Bioinformatics Core, Interdisciplinary Center for Biotechnology Research, University of Florida; George Washington University, Washington, D.C

## Abstract

Melanomas are heterogeneous and adopt multiple transcriptional states that can confer an invasive phenotype and resistance to therapy. Little is known about the epigenetic drivers of these cell states, limiting our ability to regulate melanoma heterogeneity and tumor progression. Here we identify stress-induced HDAC8 activity as the driver of a neural crest stem cell (NCSC)-like transcriptional state that increased the formation of melanoma brain metastases (MBM). Exposure of melanocytes and melanoma cells to multiple different stresses led to HDAC8 activation, a switch to a NCSC gene expression signature and the adoption of an amoeboid, invasive phenotype. This cell state enhanced the survival of melanoma cells under shear stress conditions and increased the formation of metastases in the brain. scRNA-seq analyses showed that HDAC8 expression was correlated with the NCSC cell state in clinical MBM specimens. ATAC-Seq and ChIP-Seq analysis showed HDAC8 to alter chromatin structure by increasing H3K27ac and accessibility at c-Jun binding sites without changing global histone acetylation. The increased accessibility of Jun binding sites was paralleled by decreased H3K27ac and accessibility at MITF binding sites and loss of melanoma-lineage gene expression. Mass spectrometry-based acetylomics demonstrated that HDAC8 deacetylated the histone acetyltransferase (HAT) EP300 leading to its enzymatic inactivation. This, in turn, led to an increased binding of EP300 to Jun-transcriptional sites and decreased binding to MITF-transcriptional sites. Increased expression of EP300 decreased invasion and increased the sensitivity of melanoma cells to multiple stresses while inhibition of EP300 function increased invasion and resistance to stress. We identified HDAC8 as a novel mediator of transcriptional co-factor inactivation and chromatin accessibility that increases MBM development.

## Introduction

Cutaneous melanoma is the deadliest form of skin cancer, with a propensity to aggressively metastasize to multiple organs [1]. One common site of melanoma metastasis development is the brain, with 40-60% of patients with advanced melanoma showing evidence of CNS involvement [2, 3]. Left untreated, melanoma brain metastases (MBMs) progress rapidly, with most patients dying within 3 months [4]. At this time, little is known about the molecular drivers of MBM development. Most studies to date have focused upon the role of phosphatase and tensin homologue (PTEN) loss and hyperactivation of AKT signaling as potential drivers of MBM development [5, 6]. There is evidence that clinical specimens of MBM have reduced PTEN expression/increased AKT phosphorylation compared to melanoma metastases in other sites [7]. Additionally, studies in mouse models have shown that myristolated AKT1 on a background of PTEN loss increased the formation of MBM [8]. Although there is evidence from lung and breast cancers that brain metastasis development may be associated with the acquisition of secondary genetic mutations (e.g. ERBB2 and components of the PI3K/AKT/mTOR pathway) [9], MBMs tend to have similar mutational landscapes to the extracranial tumors [10], suggesting a possible role for epigenetic drivers.

Melanoma is a heterogeneous tumor whose constituent cancer cells adopt a range of transcriptional states. High dimensional single cell analysis has identified up to 4 melanoma cell states that are defined as 1) a de-differentiated/invasive cell state (marked by low expression of the melanocyte-lineage factor MITF, high Axl expression and invasion/adhesion genes), 2) a neural crest stem cell (NCSC)-like state (characterized by low MITF expression and increased neural-crest gene expression), 3) a melanocytic/differentiated state (e.g. expressing melanocyte lineage genes such as MITF and those involved in pigmentation), and 4) a transitory state with a gene expression profile reflecting both the neural-crest and differentiated phenotypes [11, 12]. Treatment of melanomas with targeted therapy alters the heterogeneity of the tumor leading to an expansion of melanoma cell states that are more drug resistant, de-differentiated and more invasive [13-15]. At this time, little is known about how the transcriptional states are regulated and how these individually contribute to tumor progression.

We previously identified the class I histone deacetylase HDAC8 as a driver of BRAF and BRAF-MEK inhibitor resistance in *BRAF*-mutant melanoma [16]. As HDAC8 activity also increased in response to other stresses, we reasoned that increased HDAC8 activity could be critical for melanoma transcriptional state switching. In the present study, we demonstrate that HDAC8 is a stress-induced regulator of the NCSC-like transcriptional state in melanoma cells. Phenotypic studies showed the HDAC8 drives the melanoma cells to adopt an amoeboid phenotype that promoted survival under shear stress conditions, enhanced invasive capacity and increased the formation of melanoma brain metastases. Mechanistically, HDAC8 led to 1) the deacetylation of EP300, leading to an inhibition of its enzymatic activity and a switch in its association from MITF sites to Jun sites and 2) increased H3K27 acetylation and chromatin accessibility at Jun-target gene transcriptional sites. Our work provides the first evidence that HDAC8 is a regulator of the NCSC state in melanoma that leads to increased brain metastasis.

## Materials and Methods

### Cell culture and Reagents

The FOM173, NHEM, 1205Lu, A375, M233, SK-MEL-28, WM164, WM793, and WM983A cell lines were a generous gift from Dr. Meenhard Herlyn (The Wistar Institute, Philadelphia, PA). Human umbilical vein endothelial cells (HUVEC) were acquired from ATCC (Manassas, Virginia). All melanoma and melanocyte cell lines were maintained in RPMI media (Corning Inc., Corning, NY) containing 5% BSA (Millipore Sigma, Burlington, MA) and a 1:10,000 dilution of plasmocin (InvivoGen, San Diego, CA). HUVEC cells were maintained in Endothelial Cell Growth Medium (Cell Applications Inc, San Diego, CA). Every 3 months, cells were tested for *Mycoplasma* contamination using the Plasmotest-Mycoplasma Detection Test (InvivoGen) with the last test date on 06/07/2022. Cell lines were authenticated by ATCC’s Human STR human cell line authentication service and cell lines were replaced from frozen stocks after 10 passages. HUVEC cells were discarded after 3 passages. The reagents dabrafenib, vemurafenib, trametinib, and I-CBP112 were ordered from Millipore Sigma.

### Generation of Plasmids and Transfection

HDAC8 and it’s corresponding EV control plasmid were a generous gift from Dr. Edward Seto and were described previously [17]. A pcDNA3.1-p300 plasmid was a gift from Warner Greene (Addgene plasmid # 23252). A CREBBP plasmid (NM_004380) and the corresponding EV Control plasmid was ordered from Origene (Rockland, MD). EP300 siRNA (siEP300_1: SASI_Hs01_00052818 and siRNA_2: EHU155151) was ordered from Millipore Sigma and non-targeting control siRNA was ordered from Thermo Fisher. For transfection, cells were placed in OPTI-MEM media in the presence of the plasmid and Lipofectamine 2000 (Thermo Fisher Scientific, Waltham, MA). Transfection was achieved following the manufacturer’s protocol. Transfected HDAC8 and EV cells were maintained in hygromycin (Thermo Fisher Scientific). EP300 and CREBBP expressing cells were maintained in G418 sulfate (Millipore Sigma).

### Western Blotting and Immunoprecipitation

Lysates were acquired and Western Blots were performed as previously described [16]. The anti-HDAC8 antibody was described in [17]. Antibodies against SMC3 (# 5696, D47B5), p-EGFR (Y1068, # 3777, D7A5), EGFR (#4267, D38B1), p-c-Jun (S73, #3270, D47G9), c-Jun (# 9165, 60A8), HDAC1 (#34589, D5C6U), HDAC2 (#57156, D6S5P), HDAC3 (#85057, D2O1K), acetyl-histone 3 (Lys27, #8173, D5E4), histone3 (#4499, D1H2), EP300 (# 86377, D8Z4E), and CREBBP (# 7389, D6C5) were purchased from Cell Signaling Technologies (Danvers, MA). An anti-acetyl-SMC3 antibody was a kind gift from Forma Therapeutics. An Anti-MITF (NB110-10872) antibody was ordered from Novus (Littleton, CO). Anti-vinculin (V9131), anti-GAPDH (G9545) antibodies were ordered from Millipore Sigma.

Lysates for immunoprecipitation assays were collected and assayed as previously described [16]. Briefly, 500 µg of protein was incubated with an anti-EP300 antibody (CST, # 86377, D8Z4E,1;100 dilution) or an IgG control (CST, #3900, DA1E, 1:100) under constant rotation overnight at 4 degrees. Lysates were incubated with Protein A/G agarose (Thermo Fisher Scientific) for 4 hours followed by isolation of beads by centrifugation. The resulting protein/bead complexes were disassociated with loading buffer before being collected and probed with an anti-acetyl antibody (CST, #9441) by Western Blot. Western blot and immunoprecipitation protein levels were quantified using ImageJ Software. All Western Blots were normalized to loading controls for analysis and immunoprecipitations were normalized to input controls.

### Cell Viability Assays

Cells were treated with drug for 72 hours, collected and stained with Annexin V APC (BD biosciences, Franklin Lakes, NJ) and propidium iodide (PI, Millipore Sigma). Fluorescence was read on a FACSCanto (BD biosciences) and apoptotic cells were measured for Annexin V positive/PI negative populations using FloJo software (v10.7.1).

For cell death following UV irradiation and hypoxia, 300 cells were analyzed for cell death using a 0.4% trypan blue solution (Thermo Fisher Scientific) 24 hours after stress induction. Hypoxic conditions were maintained in a Hypoxia Incubator Chamber (STEMCELL Technologies, Vancouver, Canada) for 24 hours at 94% N_2_, 1% O_2_ and 5% CO_2_. Cells were UV-irradiated for 15 minutes (13.85 KJ/m^2^) using a UV Solar Simulator UV SOL (Newport Technologies, Irvine, CA) with the UV generated from the simulator consisting of 8.0% UVB and 92.0% UVA. Cells were incubated for 24 hours and cell death was measured.

For long term viability following continuous vemurafenib exposure, a colony formation assay was performed. Briefly, 1,000 cells were placed in a 6 well plate. Cells were treated with drug for 21 days, replacing drug and media twice weekly, before being stained with a 0.5% Crystal Violet solution (1:1 water to methanol) for 4 hours under constant agitation. Colonies were quantified for absorption at 580 nm following acetic acid permeabilization.

For shear stress viability, 10,000 melanoma cells were plated on a 0.4 mm channel µ-Slide I Luer (Ibidi, Martinsried, Planegg, Germany). The cells were incubated at 37 degrees Celsius for 24 hours with a constant 10 dyne/cm^2^ unidirectional flow using the Ibidi perfusion pump system. Cells were subsequently stained with Calcein AM (Molecular Probes, Eugene, OR) and PI. Cells were imaged using an EVOS 5100 autofluorescence imager (Thermo Fisher Scientific).

### Adhesion/Migration assays

For 3D spheroid assays, 10,000 cells were incubated on agar for 72 hours to form spheroids. Cells were then transferred to a Type 1 Bovine Collagen matrix (Advanced Biomatrix, Carlsbad, CA) as previously described [18]. Cells migrated for 36-48 hours and images were taken using a Nikon Eclipse Ci-L fluorescent microscope (Tokyo, Japan).

For transendothelial migration assays, HUVEC cells were plated and grown to confluence in transwell inserts as previously described [18]. 10,000 DiI-stained cells were allowed to migrate through the HUVEC coated membrane for 1 hour. Non-migratory cells were removed followed by imaging with an EVOS 5100 autofluorescence imager.

For collagen adhesion assays, 20,000 cells were grown on a Type I Bovine Collagen/PBS (pH 7.0) matrix for 48 hours. Cells were subsequently imaged using a Nikon Eclipse Ci-L fluorescent microscope.

### HAT activity assay

To measure EP300 activity, a Histone Acetyltransferase Activity Assay Kit (cat. ab239713, Abcam, Cambridge, United Kingdom) was used following the manufacturers protocol. For collection of the lysates, 500 µg of protein was immunoprecipitated with an anti-EP300 or IgG control. Fluorescence was read (Ex/Em=535/587) at 35 and 40 minutes and data were analyzed following manufacturers protocol.

### RNA Sequencing and Data Analysis

Total RNA was extracted with the Direct-zol™ RNA MiniPrep Plus Kit (Zymo Research, Irvine, CA). Library preparation and sequencing were performed by Novogene using TruSeq RNA Library Preparation Kit and NovaSeq 6000 (Illumina, San Diego, CA) with PE150. Short reads were filtered and trimmed using Trimmomatic (v 0.36). QC on the original and trimmed reads was performed using FastQC (v 0.11.4) and MultiQC (v 1.1). The reads were aligned to the transcriptome using STAR (v 2.7.3a). Transcript abundance was quantified using RSEM (v 1.2.31). Differential expression analysis was performed using DESeq2, with an FDR-corrected *P*-value threshold of 0.05 and a log_2_fold change of +/- 0.58. Gene ontology analysis of biological processes was performed using Enrichr and Gene set enrichment analysis (GSEA).

### ChIPmentation Sequencing and Data Analysis

Chromatin immunoprecipitation coupled with next-generation sequencing (ChIP-Seq) was performed with a reference exogenous genome in combination with Tagmentation for library preparation as previously described [19]. Cell lines were cross-linked with 0.8% formaldehyde and quenched with 0.125M glycine. Cells were lysed (10% glycerol, 1% Triton TX-100, 0.1 mM EDTA, 14mM NaCl 25mM Hepes pH 7.5), washed and 10×10^6^ cell nuclei were resuspended in 1 ml sonication buffer (1 mM EDTA, 10mM Tris-HCl pH 7.5). Nuclei were sonicated using a Covaris E220 Focused-ultrasonicator (Covaris Inc., Woburn, MA) for 30 minutes (15% Duty Cycle, 200 cycles/burst, and Peak incidence 140). After pre-clearing with Protein A/G Dynabeads (Invitrogen), DNA were immunoprecipitated using antibodies including anti-H3K27ac (Active Motif, #39133, Carlsbad, CA), anti-H3K27me3 (CST, #9733), and IgG (CST, #3900) overnight at 4°C. Precipitated chromatin was washed and incubated in Tagmentation buffer containing Tn5 transposase enzyme (Illumina). Samples were reverse crosslinked for 4 hours at 65°C, followed by DNA purification and amplification for the library with index primers using KAPA HiFi HotStart Kit (Roche, Indianapolis, IN). The libraries were purified, size-selected for between 200 bp and 700 bp using Ampure beads XP (Beckman Coulter, Brea, CA), and pooled for Novaseq 6000 S4 2×150 flow cell (ICBR Next-Gen Sequencing Core, University of Florida). The input sequences were trimmed using Trimmomatic. Quality control was performed before and after trimming using FastQC. The input sequences were then aligned to the GRCh38 genome using Bowtie (v 2.3.5.1). Peak detection was performed using MACS (v 2.1.2) and motif finding on peak regions was performed with the HOMER find-Motif’s function.

### ATAC Sequencing and Data Analysis

Nuclei were isolated from 200,000 cells and ATAC-Seq was performed on aliquots of 50,000 nuclei as previously described [20]. ATAC-seq libraries were purified by size selection using Ampure XP beads (Beckman Coulter, Brea, CA), and library quality assessed by fragment analyzer and qPCR for GAPDH and a genomic desert region. Sequencing of pooled libraries was performed on a Novaseq 6000 S4 2×150 flow cell (ICBR Next-Gen Sequencing Core, University of Florida). Reads were trimmed using Trimmomatic (v 0.36), and QC on the original and trimmed reads was performed using FastQC (v 0.11.4) and MultiQC (v 1.1). The reads were aligned to the human genome version GRCh38 using Bowtie (v. 2.3.3), and ATAC peak calling was performed using the MACS (v 2.1.2). Differential peak analysis was performed using DASA (https://github.com/uf-icbr-bioinformatics/dasa). Custom scripts were used to produce heatmaps and enrichment plots.

### ChIP assay

ChIP was adapted from the manufacturers protocol (Rockland, Pottstown, PA). 20 million cells were cross-linked with 1.0% formaldehyde (30 minutes) and quenched with 0.125M glycine. Cells were Dounce homogenized and extracted chromatin were sonicated using a pulse sonicator (Thermo Fisher Scientific, 15 seconds at 50% on, 1 minute off, 12 times). Samples were pre-cleared with Protein A/G Agarose and a 1/20 aliquot was saved as input control. DNA were immunoprecipitated using an anti-EP300 (ab14984, Abcam) and anti-IgG2a (ab18413, Abcam) antibody overnight at 4°C. Following a 4-hour protein A/G incubation, samples were de-crosslinked at 65 degrees C for 5 hours.

DNA was purified and subsequently amplified using the following primers:

Axl: 5’ GCTCACATTCTCAGCATTCC 3’

3’ GGAAGACCTGGTGAGCATGC 5’

MLANA: 5’ GCTGTAACAGAGCTGTAACAGAGC 3’

3’ GGAGTTGTTGAATCAGCTGCC 5’

DNA was run on a 7900HT Fast Real-Time PCR System (Thermo Fisher Scientific) for 40 cycles using TaqMan master mix (Applied Biosystems, Waltham, MA). Samples were normalized to 5% input control.

### scRNA-Seq data analysis

To investigate the relationship between the NCSC signature and HDAC8 expression in tumor cells, we performed principal component analysis (PCA) of the neural crest stem cell-like signature [11] using single cell RNA-seq (scRNA-Seq) data generated from 9,862 tumor cells by 10X genomics platform from 43 melanoma patient specimens [21].The first principal component score (PC1 score) is used as surrogate of this gene signature. The tSNE projection was used to visualize the HDAC8 gene expression and PC1 score of the Neural crest stem cell-like signature at single cell level. HDAC8 mean expression (in log2 normalized scale) is used to summarize the HDAC level for each patient sample. A Spearman correlation analysis was performed to investigate the correlation between the PC1 score and mean HDAC8 expression of these 43 patient samples.

### Acetylomics

Cells were lysed in denaturing lysis buffer containing 8M urea, 20 mM HEPES (pH 8), 1 mM sodium orthovanadate, 2.5 mM sodium pyrophosphate and 1 mM β-glycerophosphate. The proteins were reduced with 4.5 mM DTT and alkylated with 10 mM iodoacetamide. Trypsin digestion was carried out at room temperature overnight, and tryptic peptides were then acidified with 1% trifluoroacetic acid (TFA) and desalted with C18 Sep-Pak cartridges according to the manufacturer’s procedure (WAT051910, Waters Corp, Milford, MA). The peptides were then frozen on dry ice before lyophilization. Following lyophilization, the dried peptide pellet was re-dissolved in IAP buffer containing 50 mM MOPS pH 7.2, 10 mM sodium phosphate and 50 mM sodium chloride. Acetyl lysine-containing peptides were immunoprecipitated with immobilized anti-Acetyl-Lysine Motif (Ac-K) antibody (#13416, CST). The acetyl-lysine peptides were eluted twice with 0.15% TFA, and the volume was reduced to 20 µl via vacuum centrifugation. A nanoflow ultra-high performance liquid chromatograph (RSLC, Dionex, Sunnyvale, CA) interfaced with an electrospray bench top quadrupole-orbitrap mass spectrometer (Q Exactive Plus, Thermo Fisher Scientific) was used for liquid chromatography tandem mass spectrometry (LC-MS/MS) peptide sequencing experiments. The sample was first loaded onto a pre-column (C18 PepMap100, 100 µm ID x 2 cm length packed with C18 reversed-phase resin, 5 µm particle size, 100 Å pore size) and washed for 8 minutes with aqueous 2% acetonitrile containing 0.04% trifluoroacetic acid. The trapped peptides were eluted onto the analytical column, (C18 PepMap100, 75 µm ID x 25 cm length, 2 µm particle size, 100 Å pore size, Thermo Fisher Scientific). The 120-minute gradient was programmed as: 95% solvent A (aqueous 2% acetonitrile + 0.1% formic acid) for 8 minutes, solvent B (90% acetonitrile + 0.1% formic acid) from 5% to 38.5% in 90 minutes, then solvent B from 50% to 90% B in 7 minutes and held at 90% for 5 minutes, followed by solvent B from 90% to 5% in 1 minute and re-equilibration for 10 minutes. The flow rate on analytical column was 300 nl/min. Spray voltage was 1,900 v. Capillary temperature was set at 275 °C and RF was set at 50. Sixteen tandem mass spectra were collected in a data-dependent manner following each survey scan using 15 second exclusion for previously sampled peptide peaks. MS and MS/MS resolutions were set at 60,000 and 35,000, respectively. Database searches were performed with Mascot (Matrix Science, Boston, MA) and MaxQuant. The raw data were normalized using IRON processing. Values were converted to log(2) form with significant changes in acetylated proteins determined to be greater than the average of the data set +/- 2 standard deviations. Gene ontology of significantly deacetylated proteins were carried out using the STRING protein-protein interaction network.

### *In vivo* assays

1 million WM164_HDAC8 or WM164_EV expressing cells were introduced into NOD.CB17-Prkdcscid/J mice by intracardiac injection. Tumors were allowed to migrate to the brain for 14 days and mice were subsequently euthanized. After euthanasia, brains and livers were collected and fixed in formalin for 24 hours followed by storage in 70 % ethanol. The collected organs with paraffin embedded. Sections were H&E stained and subjected to immunohistochemistry (IHC) with an anti-SOX10 (ab180862, Abcam, 1:100) antibody.

### Patient database analysis

Patient samples containing RNA expression data in the TCGA PanCancer Atlas melanoma dataset were interrogated for gene expression of HDAC8 and EP300. For the HDAC8 survival analysis, a z-score of +1.2 mRNA expression of HDAC8 was compared to all samples harboring a BRAF V600 mutation. For the HDAC8/EP300 analysis, a z-score of +1 was used for HDAC8 in addition to a z-score of -1 used for EP300 compared to all samples.

### Statistical analysis

For stress assays, HAT activity assay, and ChIP assay, a 1-way ANOVA was performed followed by a post-hoc students t-test. with non-significance equaling p>0.05 and *=p<0.05, **=p<0.01. For migration assays, a student’s t-test was used. Significance was determined with non-significance equaling p>0.05 and significance equaling *=p<0.05, **=p<0.01, and ***=p,0.005. For TCGA analyses, log-ranked tests were performed with significance equaling p>0.05 and significance equaling *=p<0.05. Significance in *in vivo* assays was determined by a chi-squared test on an n of 20 mice with ***=p<0.005.

## Results

### HDAC8 activity is increased following exposure of melanoma cells to stress

We began by determining if HDAC8 was an epigenetic driver of cell state changes in melanoma. Western Blot profiling of a panel of melanoma cell lines demonstrated that higher expression of the melanocyte lineage transcription factor MITF was correlated with lower HDAC8 expression and reduced levels of signaling molecules known to be regulated by HDAC8 including acetylated SMC3, EGFR (both total and phospho) and c-Jun (both total and phospho) [16, 22]. The cell lines were ranked according to their baseline sensitivity to the BRAF inhibitor vemurafenib [14] and demonstrated a link between higher MITF expression, lower HDAC8 expression and increased BRAFi sensitivity (Figure 1A). Stable expression of HDAC8 in a BRAFi sensitive melanoma cell line significantly shifted the IC_50_ to the BRAFi vemurafenib and exhibited a lower proliferation rate (Supplemental Figure 1). Likewise, introduction of HDAC8 into melanoma cell lines with low endogenous HDAC8 expression increased phospho-Jun and EGFR levels (Figure 1B).

**Figure 1:**
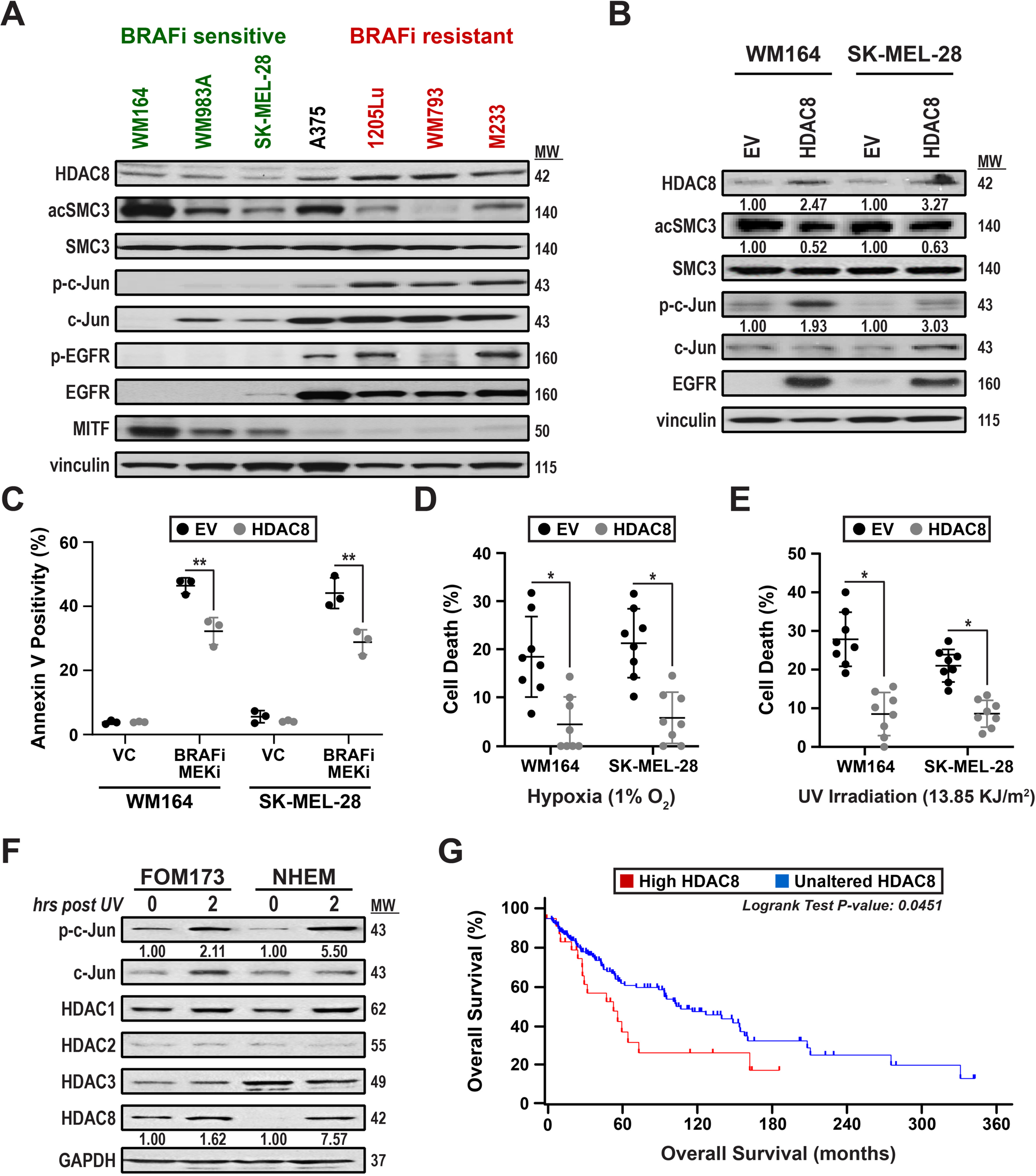
HDAC8 expression confers stress resistance in melanoma. **A:** BRAFi resistant cells display increased HDAC8, c-Jun, and EGFR protein levels with decreased MITF. Cell lines treated with vemurafenib (BRAFi) were organized into BRAFi sensitive (IC50 < 1 µM; WM164, WM983A, and SK-MEL-28 cells), BRAFi intermediate (IC50=1; A375), and BRAFi resistant (IC50 >1 µM; 1205Lu, WM793, and M233 cells) cells. Cell lines were probed for HDAC8, acetylated SMC3 (acSMC3), phospho-EGFR (p-EGFR), EGFR, phospho-c-Jun (p-c-Jun), c-Jun, and MITF levels by Western blot. **B:** Forced expression of HDAC8 increases activated c-Jun and EGFR protein expression. Cells transfected with an empty vector (EV) or HDAC8 construct were probed for HDAC8, acSMC3, SMC3, p-c-Jun, c-Jun, and EGFR levels by Western blot. Levels of HDAC8, acSMC3, and p-c-Jun were quantified by ImageJ. **C:** HDAC8 confers resistance to BRAF-MEK inhibitor resistance. Cells were treated with vehicle control (1:1000 DMSO, VC), or with the BRAFi dabrafenib and the MEKi trametinib (100 nmol/L dabrafenib, 10 nmol/L trametinib, BRAFi-MEKi) for 72 hours. Apoptosis was measured by Annexin V/PI staining using flow cytometry. **D** and **E:** HDAC8 expression mediates resistance to cellular stress. Cells were treated with **(D)** 1% O_2_ for 24 hours or **(E)** 13.85 KJ/m^2^ UV irradiation followed by a 24-hour incubation. After treatment, cell death was measured by trypan blue inclusion. **F:** HDAC8 and phospho-c-Jun are increased after induction of stress in normal melanocytes. Cell lines were treated with 13.85 KJ/m^2^ UV irradiation for indicated time points. After treatment, cells were collected and probed for p-c-Jun, c-Jun, HDAC1, HDAC2, HDAC3, and HDAC8 by western blot. Levels of HDAC8 and p-c-Jun were quantified by ImageJ. **G:** Altered HDAC8 expression was associated with a decrease in patient survival. Patient samples with altered HDAC8 mRNA expression levels (+ 2-fold expression) were compared to unaltered patient samples using the TGCA PanCancer Atlas melanoma database. Significance was determined by a log rank test. Significance in **(C), (D)**, and **(E)** was determined by a one-way ANOVA followed by a post hoc t-test with *=p<0.05, **=p<0.01, and #=p>0.05. Experiments were run 3 independent times with an n of 3 in each cohort in **(C)** and an n of 8 in each cohort in **(D)** and **(E)**.

As transcriptional switching could be a possible response to exogenous stress, we next asked whether introduction of HDAC8 increased the survival of melanocytic cells under different microenvironmental conditions. Increased HDAC8 expression led to increased survival of melanoma cells exposed to hypoxia, UV irradiation and after treatment with BRAF-MEKi therapy (Figures 1C-E). To determine if this was a conserved response between melanocytes and melanoma cells, we treated 2 primary human melanocyte cell lines with UV irradiation and noted increased expression of HDAC8 and increased phosphorylated c-Jun expression (Figure 1F). The clinical relevance of increased HDAC8 expression was confirmed through interrogation of the TCGA Skin Cutaneous Melanoma database which showed that increased HDAC8 expression (n=25) was predictive of poorer overall survival compared to patients with unaltered HDAC8 levels (n=130) (Figure 1G).

### HDAC8 drives a switch to an invasive neural crest stem cell-like melanoma state

RNA-seq analyses were performed to determine how upregulation of HDAC8 reshaped the transcriptional landscape. It was found that 810 genes significantly increased, and 809 genes significantly decreased upon HDAC8 expression (Figure 2A). A Kyoto Encyclopedia of Genes and Genomics (KEGG) pathway analysis revealed that HDAC8 expression increased enrichment for pathways important for melanoma migration including Focal Adhesion, PI3K-AKT signaling, ECM-Receptor Interactions, Regulation of Actin Cytoskeleton, MAPK signaling and RAP1 signaling (Figure 2B) while decreasing enrichment of Metabolic Pathways, Central Carbon Metabolism and Oxidative Phosphorylation (Supplemental Figure 2A). An additional Molecular Signature Database (MSigDB) analysis focused on Hallmark genesets showed that HDAC8 expression increased genes associated with an epithelial-to-mesenchymal-transition (EMT) while decreasing genes involved in the Reactive Oxygen Species Pathway and Heme Metabolism (Supplemental Figures 2B-C). As these data suggested the HDAC8-driven transcriptional program was associated with an invasive, metastatic phenotype we next determined whether increased HDAC8 expression also correlated with previously identified melanoma transcriptional states.

**Figure 2:**
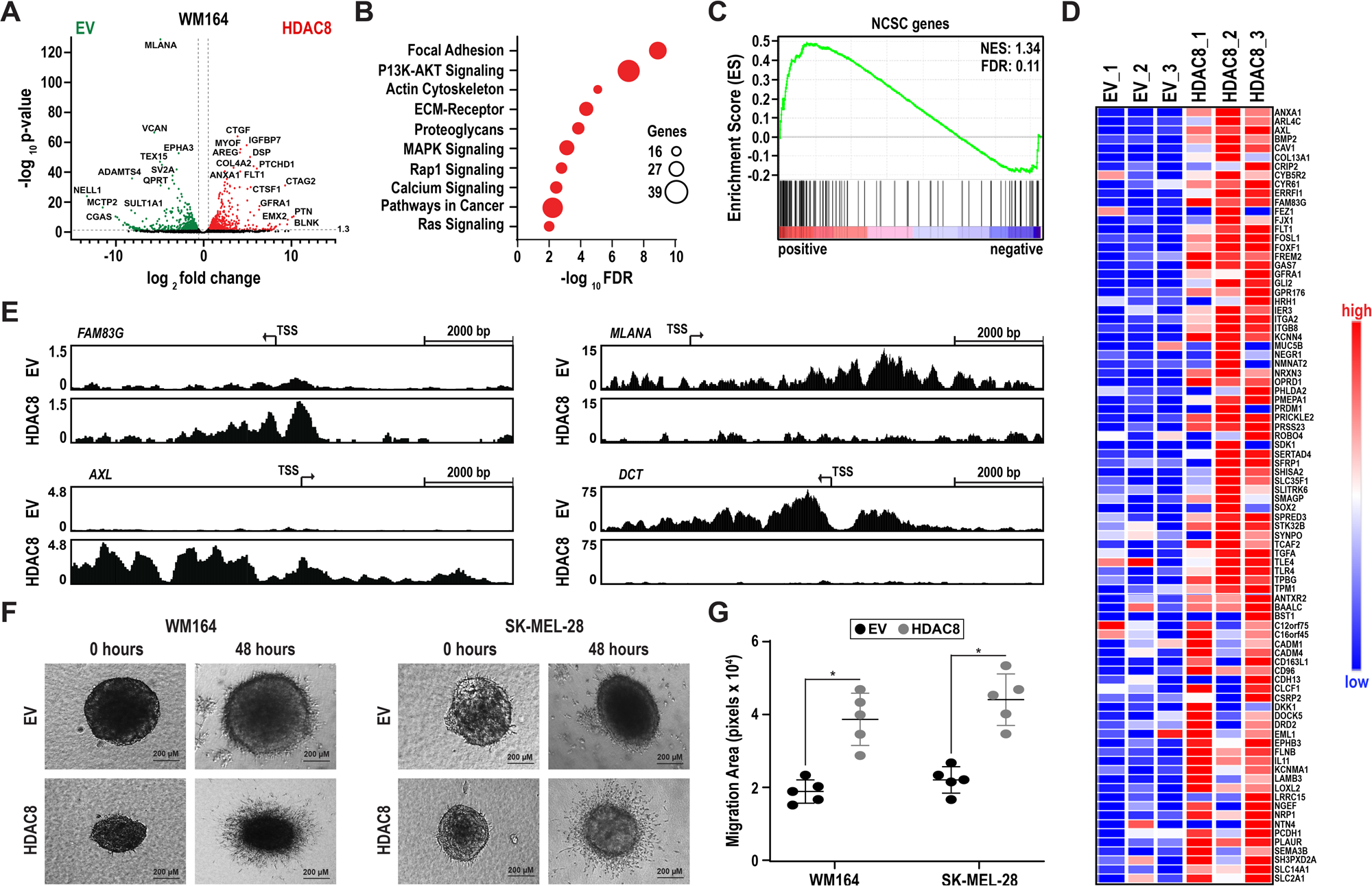
HDAC8 drives an invasive neural crest stem cell-like melanoma state. **A:** HDAC8 induces significant global changes in gene expression. RNA-seq was performed on EV and HDAC8 expressing WM164 cell lines with significantly changed genes assigned a log_2_ fold change value +/- 0.58 and a -log_10_ p-value above 1.3. **B:** Expression of HDAC8 enriches for pathways involved in adhesion and migration. A KEGG pathway analysis was performed on significantly changed genes using RNA-seq data. **C-D:** Introduction of HDAC8 increased expression of genes associated with a neural crest stem cell-like state (NCSC). **C:** A Ranked Order analysis was performed on NCSC and NCSC/undifferentiated genes using GSEA software. **D:** Heat maps were made of significantly altered NCSC genes comparing EV and HDAC8 expressing cells. **E:** Increased initiation of transcription in NCSC genes (*FAM83G* and *AXL*) upon expression of HDAC8 and decreased initiation of transcription in melanocytic genes (*MLANA* and *DCT*) upon expression of HDAC8. ChIP-Seq was performed on EV and HDAC8 expressing WM164 cells using an acetyl-H3K27 antibody followed by interrogation of gene promoter regions. **F-G:** Expression of HDAC8 increases invasion in melanoma cells. **F:** EV and HDAC8 expressing melanoma spheroids were plated in collagen for indicated time points. **G:** Spheroid invasion area was quantified using ImageJ software. Significance was determined using a student’s t-test with * = p<0.05. Experiments were run 3 independent times with an n of 5 in each cohort.

We determined that HDAC8 expression significantly increased genes associated with the invasive, drug resistant, undifferentiated phenotype (19 of 53 genes changed, NES score: 1.15, FDR: 0.17) characterized by Hoek and colleagues [23] while showing significant decreases in genes associated with the proliferative, drug-sensitive, differentiated phenotype (33 of 48 genes changed, NES score: -3.03, FDR <0.001) (Supplemental Figures 3A-B). More recent high-dimensional analyses of melanoma heterogeneity identified at least 4 different cell states, including an undifferentiated embryonic stem cell (ESC) state, a neural crest stem cell (NCSC)-like state, a transitory state, and a differentiated melanocyte state [11]. Analysis of our dataset demonstrated that HDAC8 expression enriched for genes associated with the NCSC state, whereas genes associated with the differentiated melanocyte state were significantly downregulated (Figures 2C-D, Supplemental Figure 4A). Surprisingly, genes associated with the undifferentiated ESC state were not enriched upon expression of HDAC8, indicating a specific NCSC phenotype regulated by HDAC8 (Supplemental Figure 4B).

We next verified increased initiation of transcriptional activity of NCSC genes by analyzing H3K27ac ChIP-Seq data in the promoter regions of NCSC genes in WM164 and 1205Lu cells. Genes associated with the NCSC state, including *AXL, FAM83G, FOSL1*, and *SOX2* had increased H3K27ac activity in HDAC8 expressing cells (Figure 2E) while genes associated with a melanocytic state, including *MLANA, DCT, TYR*, and *PMEL* had decreased H3K27ac activity in HDAC8 expressing cells (Figure 2E, Supplemental Figure 5). As many genes that were upregulated in our RNA-seq and ChIP-Seq analysis are linked to increased invasion in melanoma, we performed 3D spheroid collagen invasion assays and found that expression of HDAC8 led to increased cell invasion when compared to EV cells (Figures 2F-G).

### The HDAC8-driven NCSC transcriptional state leads to the adoption of an invasive, shear stress-tolerant, amoeboid phenotype and increased brain metastasis

Cells that metastasize to the brain must overcome multiple obstacles including survival under the high shear stress conditions of the circulatory system, extravasation through the blood-brain-barrier and growth in the protease-rich environment of the brain parenchyma [24-26]. We hypothesized that the increased invasive capacity/stress tolerance of the HDAC8-driven cell state could increase MBM development. We first asked whether HDAC8 increased the resilience of melanoma cells to shear stress through use of a flow chamber system that mimics the levels of shear experienced in the general circulation [27, 28]. Exposure of melanoma cells to high levels of fluid shear stress (10 dyne/cm^2^) over 24 hr led to high levels of cell death in the control (EV) cells that were reduced by 80-90% in HDAC8-expressing melanoma cells (Figures 3A-B). Many of the genes upregulated by HDAC8 were involved in cytoskeletal rearrangement, including NCSC genes such as *SOX2* and *PROM1* that are associated with an amoeboid phenotype [29]. In line with this observation, HDAC8 expressing melanoma cells adopted a rounded, amoeboid phenotype after being plated onto collagen (Figure 3C). As the amoeboid phenotype can allow cells to squeeze through tight spaces, such as those associated with exiting blood vessels and crossing the blood brain barrier, we next performed transendothelial cell migration assays and found that HDAC8 expression led to a ten-fold increase in transmigration of melanoma cells through confluent endothelial cell layers (Figures 3D-E).

**Figure 3:**
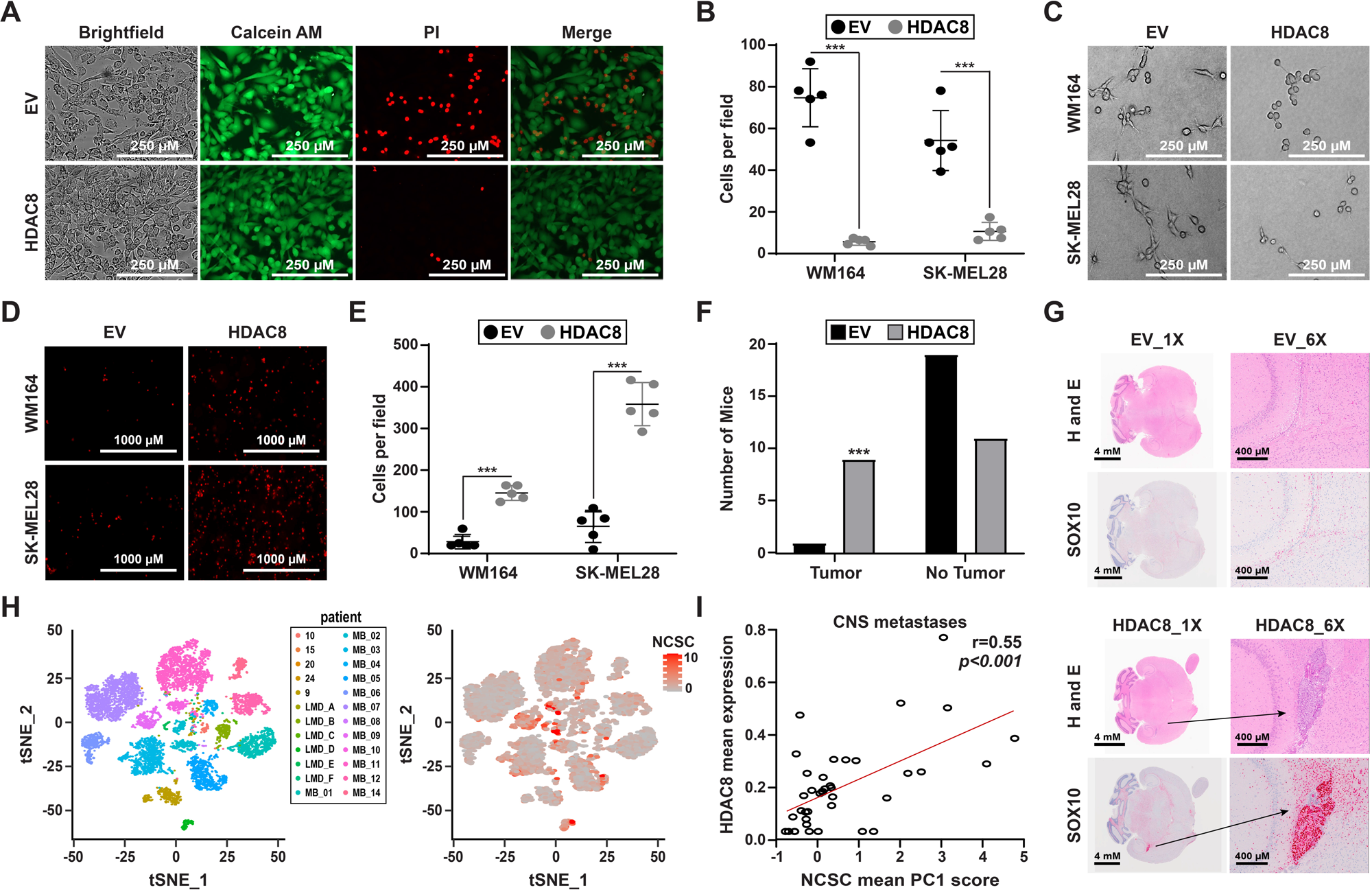
HDAC8 increases establishment of melanoma brain metastases. **A-B:** HDAC8 expression confers resistance to shear stress. Cells were incubated for 24 hours under shear stress conditions of 10 dyne/cm^2^. Cells were stained with Calcein AM/PI and imaged for cell viability. Viability was quantified as the number of PI positive cells per image. **C-E:** HDAC8 expressing cells adopt an amoeboid phenotype. **C:** HDAC8 and EV expressing cells were plated on collagen overnight followed by brightfield image collection. **D:** DiI-stained HDAC8 and EV cells were allowed to migrate through a HUVEC coated transwell membrane for 1 hour followed by imaging with fluorescence microscopy. **E:** DiI-stained cells were quantified by counting the number of cells per image. **F-G:** Introduction of HDAC8 increases metastasis to the brain *in vivo*. HDAC8 and EV expressing cells were introduced into NOD.CB17-*Prkdc*^scid^/J mice by intracardiac injection. Tumors were allowed to establish for 14 days. **F:** Number of tumors established in the brain for each condition. **G:** Embedded brain sections were stained with H&E and an IHC for SOX10 to determine tumor formation. **H:** Patient-derived scRNA-Seq data (#’s=melanoma cutaneous samples, LMD: melanoma CSF samples, MB: melanoma brain metastases samples) was interrogated for expression of NCSC genes. **I:** A Spearman correlation analysis was run to determine the relationship of HDAC8 expression and a NCSC phenotype in the scRNA-Seq database. A p-value p<0.05 corresponds to a significant correlation. Significance in **(B)** and **(E)** was determined by a student’s t-test with ***=p<0.005. Experiments were run 3 independent times with an n of 5 in each cohort. Significance in **(F)** was determined by a chi-squared test on an n of 20 mice with ***=p<0.005.

We next asked whether the HDAC8-driven NCSC phenotype altered the patterns of metastatic seeding *in vivo*. Control or HDAC8 expressing melanoma cells were injected into the left ventricle and the mice left for 2 weeks for tumors to establish. HDAC8 expressing melanoma cells showed a significantly marked increase in metastasis to the brain (9/20 mice) compared to the control (EV) cells (1/20 mice) (Figures 3F-G). Most of the HDAC8 expressing melanoma cells infiltrated the brain via the choroid plexus (5/9) (Figure 3G, Supplemental Figures 6A-C). By contrast, levels of metastasis to the liver were similar for both HDAC8 expressing cells (14/20 mice) and control (EV) melanoma cells (12/20 mice) (Supplemental Figure 6D). To address the clinical relevance of these findings we interrogated our recent single cell RNA-seq analysis of clinical MBM samples [21]. These analyses showed a significant correlation between HDAC8 expression and the NCSC-like gene expression profile in the human melanoma cells (Figures 3H-I).

### HDAC8 increases chromatin accessibility and H3K27ac of Jun-targeted genes and decreases accessibility and H3K27ac of MITF-targeted genes

We next used ATAC-Seq and ChIP-Seq to determine how HDAC8 regulated the NCSC gene transcription program. Global analysis of ChIP-Seq data revealed HDAC8 expression to have little global effect on H3K27ac at either promoter or enhancer regions (Figures 4A-B), there was likewise little alteration of global acetyl-H3 histone protein levels (Figure 4C). A HOMER analysis of H3K27ac ChIP-Seq data revealed an increase in H3K27 acetylation at Jun sites when HDAC8 was introduced, whereas control cells exhibited an increase in H3K27 acetylation at transcription factors important for cell cycle progression, including B-myb and A-myb (Figure 4D). An ATAC-seq analysis of peak numbers/length in genes showed similar levels of accessibility and DNA fragment sizes in HDAC8 expressing cells and EV cells (Figure 4E). While overall accessibility was similar throughout the genome, promoters showed increased accessibility in HDAC8 expressing cells (Figure 4F) whereas little difference was seen at non-promoter sites (Figure 4G). An analysis of the motifs with increased accessibility upon HDAC8 expression included multiple AP-1 transcription factors (Fra1, ATF3, Jun) and TEADs (Figure 4H, Supplemental Figure 7). Multiple motifs also showed downregulation upon HDAC8 introduction including CTCF, multiple SOX transcription factors, and the melanocyte lineage transcription factor MITF (Figure 4I, Supplemental Figure 7).

**Figure 4:**
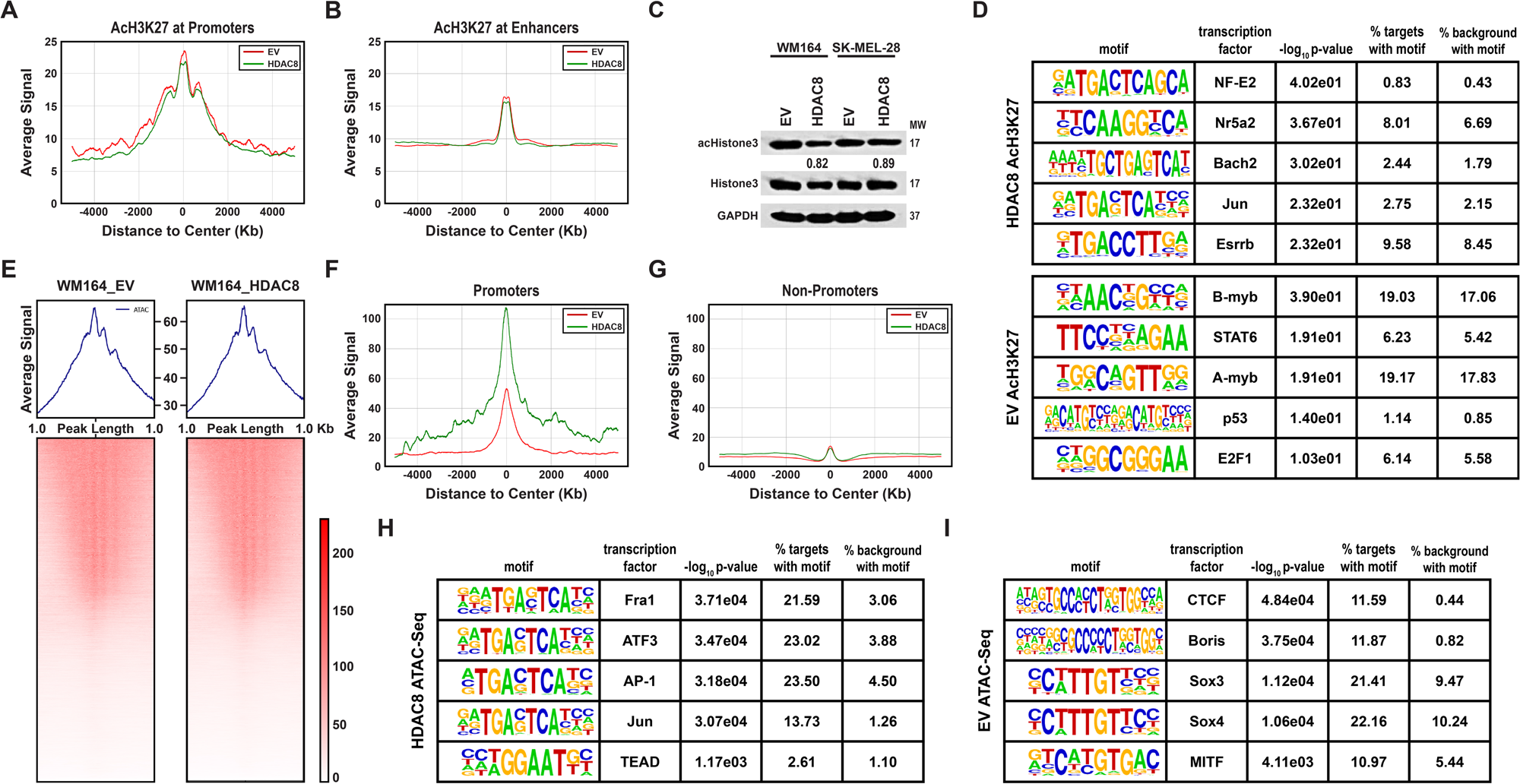
HDAC8 activation induces changes in chromatin accessibility of Jun and MITF targeted genes. **A-C:** HDAC8 expression does not alter global H3K27 acetylation. A ChIP-Seq analysis was performed on HDAC8 expressing and EV expressing melanoma cells at both **(A)** promoter and **(B)** enhancer regions. **C:** Cell lines were probed for acetyl-Histone3 andtotal-Histone 3 levels by western blot. Levels of acetyl-Histone3 were normalized to total-Histone3 and quantified by ImageJ. **D:** HDAC8 expression increases H3K27 acetylation at Jun binding motifs. A HOMER analysis was performed on HDAC8 and EV expressing cells. Shown are enriched transcription factor binding motifs in HDAC8 and EV expressing cells. **E-G:** HDAC8 expression increases accessibility in promoter regions of a specific subset of chromatin. ATAC-Seq was performed on HDAC8 expressing and EV expressing cells. **E:** A global analysis of all open chromatin was performed. A global analysis was performed on HDAC8 expressing and EV expressing melanoma cells at both **(F)** promoter and **(G)** non-promoter regions. **H-I:** HDAC8 expression increases accessibility in Jun targeted genes and decreases accessibility of MITF targeted genes. A HOMER analysis was performed on ATAC-Seq data. HDAC8 was associated with increased accessibility at **(H)** Jun binding sites while the EV control had increased accessibility at **(I)** MITF binding sites.

### HDAC8 deacetylates and inactivates EP300 leading to increased Jun transcriptional activity

Bioinformatic integration of ATAC-Seq and ChIP-Seq data suggested HDAC8 expression confers enhancement of accessibility/transcriptional activation of genes in pathways associated with extracellular matrix assembly, cell spreading and differentiation (Figure 5A, Supplemental Figure 8). Although the ATAC-Seq and ChIP-Seq data showed HDAC8 mediated few global changes in H3K27ac or chromatin accessibility, changes at discrete Jun and MITF sites were noted. Interestingly, many of the genes implicated in the NCSC phenotype, such as *SOX2* are regulated by c-Jun and demonstrated increased accessibility/H3K27 acetylation in the HDAC8 expressing cells (Figure 5B), while MITF regulated genes, such as *MLANA* (involved in the melanocytic phenotype) had decreased accessibility (Figure 5C). Of relevance to the development of MBM, we noted that HDAC8 expression led to increased H3K27ac and chromatin accessibility of multiple Serpins (e.g., Serpin E1, E2 and A1), protease inhibitors that facilitate the establishment of brain metastases [25] (Supplemental Figure 9). Together these data suggested that HDAC8 introduction led to a switch from MITF-driven transcriptional programs to those regulated by c-Jun.

**Figure 5:**
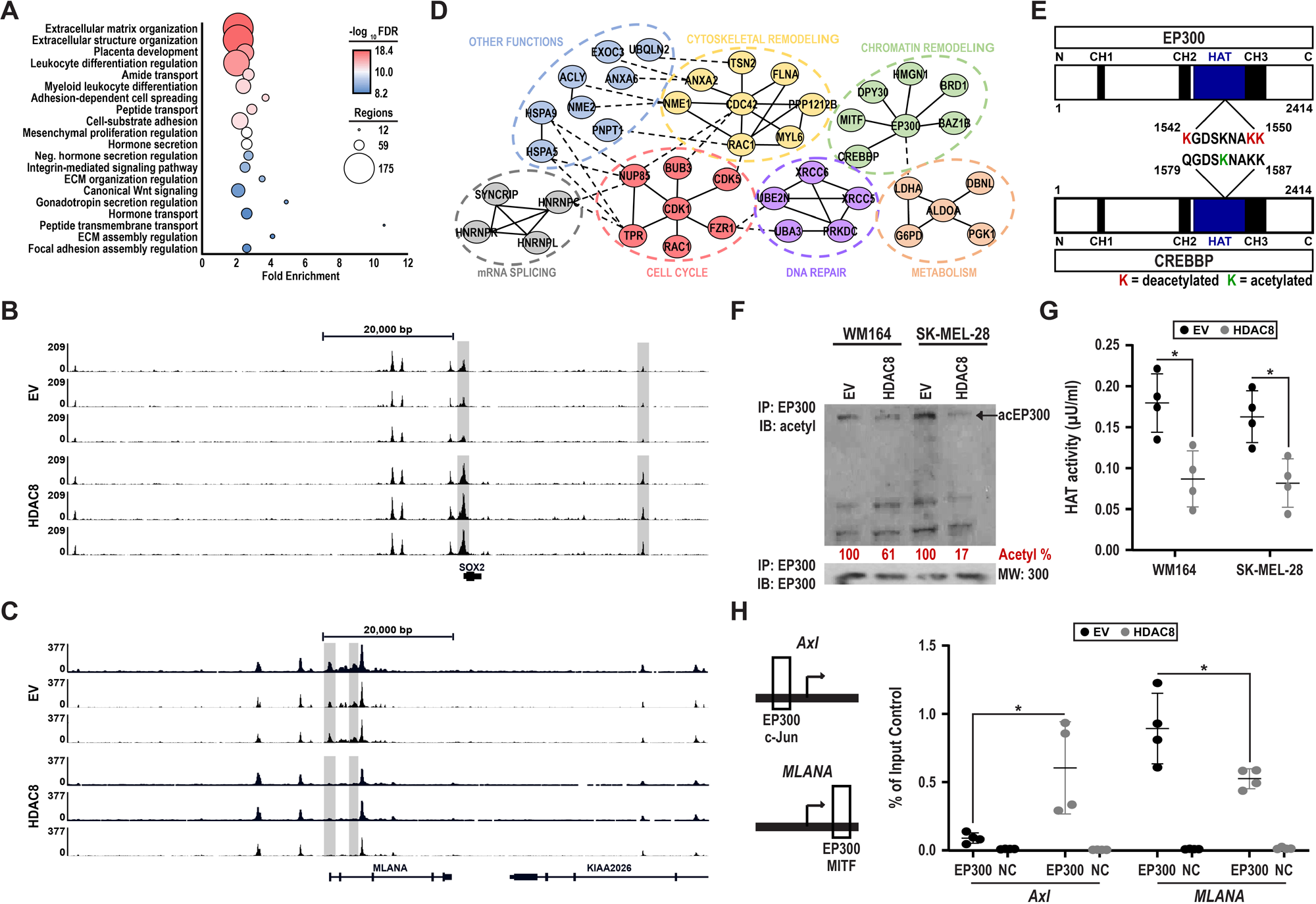
HDAC8 directly deacetylates and inactivates EP300. **A:** HDAC8 expression revealed enriched chromatin accessibility/initiation of transcription for genes involved in migration and invasion. A GREAT analysis was performed on genes with enhanced accessibility/H3K27 acetylation around their transcription start sites. **B-C:** HDAC8 expression increases accessibility in NCSC genes while EV increases accessibility in melanocyte genes. There is increased accessibility in the promoter region of the **(B)** NCSC gene SOX2 while there is decreased accessibility in the promoter region of the **(C)** melanocyte gene MLANA upon HDAC8 expression. **D-E:** The HAT co-activators EP300 and CREBBP are deacetylated upon HDAC8 expression. A mass spectrometry-based acetylomics assay was performed on HDAC8 and EV expressing cells. **D:** Proteins deacetylated by HDAC8 were organized into signaling hubs using STRING. **E:** Significantly changed lysine acetylation sites in the CH3 domain of CREBBP and EP300 are shown. **F:** HDAC8 deacetylates EP300. Immunoprecipitation assays were performed with EP300 in the cell lines indicated. Lysates were probed for acetyl-lysine levels by Western Blot. Levels of acetyl-EP300 were normalized to EP300 input controls and quantified by ImageJ. **G:** HDAC8 reduces EP300 histone activity. Immunoprecipitation assays were performed with EP300 in two independent melanoma cell lines. Collected lysates were analyzed for EP300 mediated H4 histone acetylation. **H:** HDAC8 increases binding of EP300 to Jun bound promoter regions and decreases binding of EP300 to MITF bound promoter regions. A ChIP assay was performed on EP300 binding to the promoter region of *Axl* and *MLANA*. Significance in **G** was determined by a student’s t-test with *=p<0.005. Significance in **(H)** was determined by a one-way ANOVA followed by a post hoc t-test with *=p<0.05. Experiments were run 3 independent times with an n of 4 in each cohort.

As the effects of HDAC8 on global chromatin accessibility were relatively minor we next asked whether HDAC8 mediated any of its effects through regulation of non-histone targets. A proteomics analysis was performed to determine how HDAC8 affected the “acetylome” of melanoma cells (Supplemental Figure 10A shows the workflow). Significantly deacetylated proteins, including those involved in cytoskeleton remodeling, cell cycle, DNA repair, metabolism and chromatin remodeling were identified by STRING analysis (Figure 5D). One of the central hubs in the “chromatin remodeling” network was the histone acetyltransferase (HAT) EP300/CREBBP, which is known to acetylate multiple proteins including MITF and c-Jun [30, 31]. An analysis of the structure of EP300 and CREBBP identified 3 lysine residues in the HAT domain of EP300 (Lysine’s 1542, 1549, and 1550) that were deacetylated upon expression of HDAC8 while one lysine residue in CREBBP (Lysine 1583) was significantly acetylated upon expression of HDAC8 (Figure 5E, Supplemental Figure 10B). The ability of HDAC8 to deacetylate EP300 was confirmed by immunoprecipitation of EP300 from both EV and HDAC8 expressing cells and Western blotting for total protein acetylation (Figure 5F). The deacetylation of EP300 was associated with a decrease in its HAT activity (as measured by acetylation of histone H4) in two independent melanoma cell lines (Figure 5G). Identical experiments following immunoprecipitation of CREBBP saw no decrease in CREBBP acetylation or HAT activity upon expression of HDAC8 (Supplemental Figure 11). ChIP assays for NCSC and melanocytic phenotype associated genes showed that increased expression of HDAC8 was associated with increased binding of EP300 to the Jun promoter of *AXL* and a decreased binding of EP300 to the MITF promoter of *MLANA* (Figure 5H). Together these results demonstrated that HDAC8 modulated EP300 function both through inhibition of its HAT activity and by a switching its association from MITF promoter sites to Jun promoter sites.

### Inhibition of EP300 can drive the stress-resistant NCSC phenotype in melanoma cells

Since EP300 is a direct target of HDAC8, we next explored the role of EP300 in driving the stress-resistant, invasive NCSC phenotype. We first looked at the expression levels of EP300 and CREBBP across a panel of melanoma cell lines that were either sensitive or resistant to BRAFi therapy. It was noted that the cell lines that were sensitive to inhibitor therapy had increased EP300 levels (Figure 6A). Interrogation of the TCGA Melanoma database highlighted a correlation between decreased overall survival and a concurrent decrease in the levels of EP300 and increased levels of HDAC8, (change of 2-fold expression, n=37) had significantly decreased overall survival compared to patients with unaltered EP300/HDAC8 levels (n=406) (Figure 6B). Performing the same analysis with increased levels of HDAC8 and decreased levels of CREBBP did not show a significant decrease in survival (Supplemental Figure 12A-C). Functional studies were then undertaken to determine if inhibition of EP300 (mimicking its deacetylation by HDAC8) would alter sensitivity of melanoma cells to BRAFi therapy. Co-treatment of *BRAF*-mutant melanoma cell lines with the EP300/CREBBP inhibitor I-CBP112 decreased the sensitivity of the melanoma cells to BRAFi, increasing the number of colonies formed (Figures 6C-D). Silencing of EP300 through specific siRNA had similar effects and was found to decrease the level of apoptosis induced in response to BRAFi-MEKi therapy (Figure 6E), while silencing of CREBBP by siRNA was less significant (Supplemental Figure 13). The effects were associated with increased c-Jun activity, with EP300 inhibition being found to increase levels of transcriptionally active phospho-c-Jun (Figure 6F). The converse was also true, with increased expression of EP300 being found to increase cell death in response to treatment with BRAFi-MEKi. Overexpression of CREBBP did not significantly alter cell death in response to BRAFi-MEKi treatment (Figure 6G). As the NCSC cells are highly invasive, we next examined if modulating EP300 expression would alter the invasive capacity of melanoma cells. Inhibition of EP300/CREBBP increased the invasion of melanoma cells into 3D collagen matrices (Figures 6H-I) whereas overexpression of EP300 decreased cell invasion (Figure 6J-K). Together our data suggest that HDAC8 mediated deacetylation of EP300 leads to decreased HAT activity, an increase in transcriptionally active JUN and a switch from a melanocytic to a NCSC-like program (model shown in Figure 6L).

**Figure 6:**
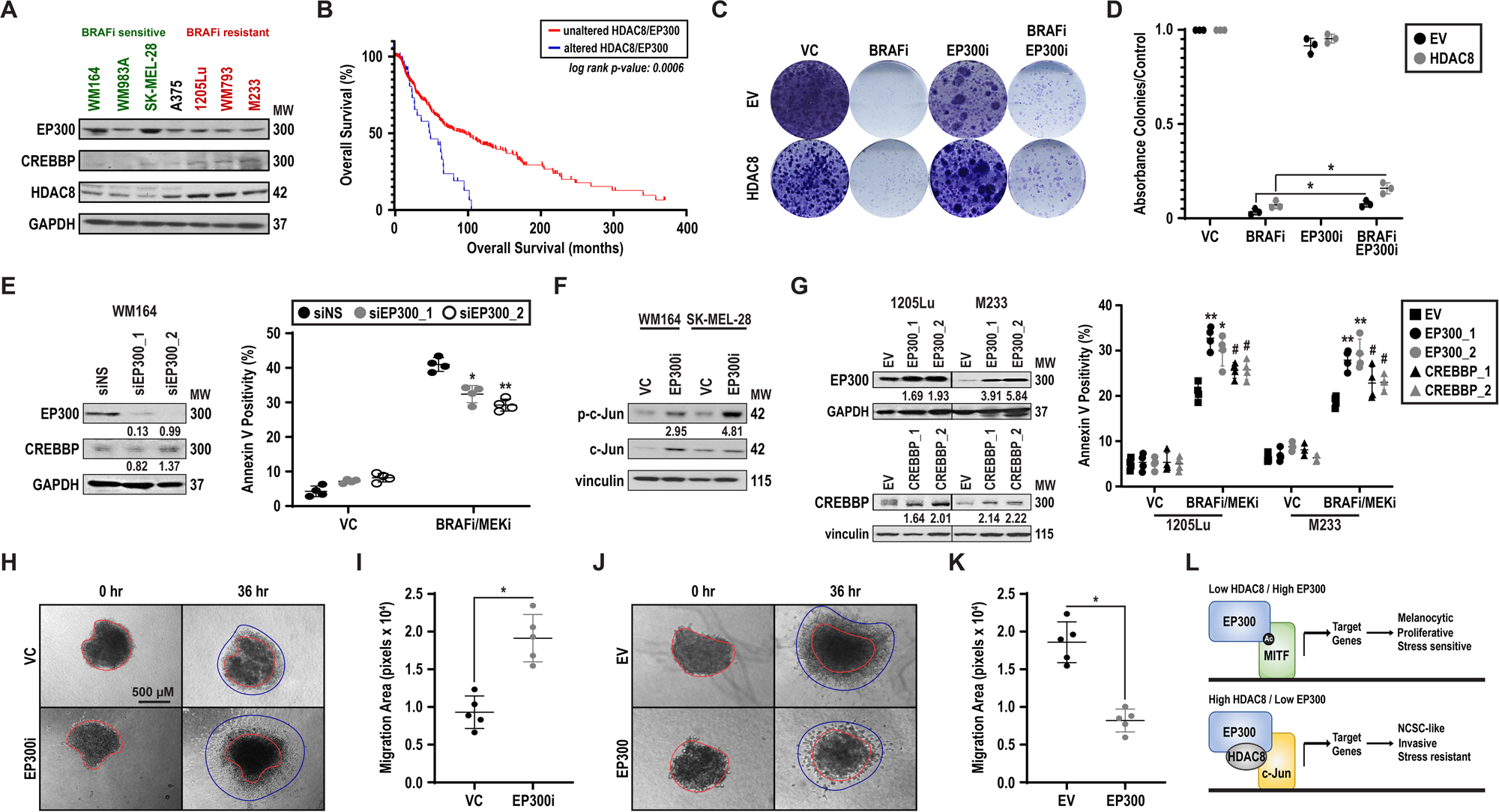
EP300 reverses HDAC8 induced stress resistance in melanoma. **A:** EP300 is downregulated in BRAFi resistant cells. Cell lines were probed for EP300, CREBBP and HDAC8 levels by western blot. **B:** Alterations in HDAC8/EP300 expression conferred a decrease in patient survival. Patient samples with decreased EP300 and increased HDAC8 RNA levels (change in 2-fold expression over all samples) were compared to unaltered patient samples using the TGCA melanoma database. **C-D:** Inhibition of EP300/CREBBP increases resistance to BRAFi. **C:** Cells were treated with vemurafenib (1 μmol/L, BRAFi), I-CBP112 (100 nmol/L, EP300i) or combination for 21 days. Colony formation was measured by Crystal Violet staining. **D:** Colonies were quantified using acetic acid permeabilization followed by ImageJ analysis. **E:** Silencing EP300 decreases sensitivity to BRAFi-MEKi. WM164 cells were transfected with a non-silencing siRNA (siNS) or an EP300 siRNA (siEP300_1 or siEP300_2). Cells were probed for EP300 and CREBBP after 72 hours transfection. Levels of CREBBP and EP300 were quantified by ImageJ. Cells were also treated with dabrafenib and trametinib (100 nmol/L dabrafenib, 10 nmol/L trametinib, BRAFi-MEKi) for 72 hours. Apoptosis was measured by Annexin V/PI staining using flow cytometry. **F:** Silencing of EP300 increases c-Jun phosphorylation. Cell lines were treated with I-CBP112 (100 nmol/L, EP300i) for 24 hours. Lysates were collected and probed for p-c-Jun and c-Jun by western blot. Levels of p-c-Jun were quantified by ImageJ. **G:** Forced expression of EP300 increases sensitivity to BRAFi-MEKi, whereas CREBBP does not. Cells with intrinsic resistance to BRAF inhibition were stably transfected with an empty vector, EP300 or CREBBP. Protein levels of EP300 and CREBBP were quantified for overexpression by ImageJ. Cells were treated with dabrafenib and trametinib (100 nmol/L dabrafenib, 10 nmol/L trametinib, BRAFi-MEKi) for 72 hours. Apoptosis was measured by Annexin V/PI staining using flow cytometry. **H-I:** Inhibition of EP300 increases migration in BRAFi sensitive cell lines. **H:** Melanoma spheroids were treated with ICBP-112 (100 nmol/L, EP300i) or VC and plated in collagen for indicated time points. **I:** Spheroid invasion area was quantified using ImageJ software. **J-K:** Introduction of EP300 decreases migration in BRAFi resistant cell lines. **J:** EP300- or EV-expressing spheroids were plated on collagen for indicated time points. **K:** Spheroid invasion area was quantified using ImageJ software. **L:** A schematic of the proposed mechanism of action for the HDAC8/EP300 modulated phenotype switch is shown. Significance in **(D), (E)**, and **(G)** was determined by a one-way ANOVA followed by a post hoc t-test with *=p<0.05, **=p<0.01, and #=p>0.05. Significance in **(I)** and **(K)** was determined by a student’s t-test with *=p<0.05. Experiments were run 3 independent times with an n of 3 in **(D)**, an n of 4 in **(E)** and **(G)**, and an n of 5 in **(I)** and **(K)**.

## Discussion

In the current study we identified a unique role for HDAC8 in driving a stress-induced neural crest phenotype in melanoma cells that increases metastasis to the brain. HDAC8 is a type I HDAC that is essential for correct embryonic development and migration of neural crest cells [32, 33] and has been implicated as a key driver of tumorigenesis in the neural crest tumor neuroblastoma in addition to other tumors such as acute myeloid leukemia and breast cancer [34-36]. HDAC8 is unique among class I HDACs, and differs from HDAC1, 2, 3 in not being phosphorylated and activated by casein kinase 2 (CK2). Instead, HDAC8 is phosphorylated by cAMP-activated PKA signaling at its N-terminus at Serine 39 [37], leading to the inhibition of its HDAC function [17]. Phosphorylation of HDAC8 also impedes its ability to translocate to the nucleus preventing it from acting on nuclear substrates such as CREB and p53 [17]. At this time, little is known about the role of HDAC8 in the transcriptional state regulation of melanocytes/melanoma cells and its potential role in melanoma progression/metastasis.

An analysis of a panel of melanoma cell lines identified those with either low or high expression of HDAC8. Cell lines with highest HDAC8 expression had evidence of phenotype switching (e.g., low MITF expression) and this was associated with *de novo* resistance to BRAFi therapy. By contrast, cell lines with the lowest HDAC8 activity had the highest MITF expression and increased BRAFi therapy sensitivity. HDAC8 expression/activity was also noted to increase in melanoma cells following exposure to multiple stresses including UV irradiation, hypoxia, and drug treatment. Previous studies have shown that HDAC8 directly deacetylates p53 under conditions of stress, leading to increased cell survival [38, 39]. Similar increases in both HDAC8 and c-Jun were also seen in melanocytes treated with UV irradiation, suggesting a conserved mechanism of stress tolerance between melanocytes and melanoma cells.

An RNA-seq analysis of the HDAC8-driven transcriptional state showed the gene expression profile was akin to the melanoma neural crest stem cell (NCSC)-like state [11]. ATAC-seq analysis demonstrated there to be increased chromatin accessibility at multiple genes that were specific to the NCSC state including *FAM83G* and *AXL* in conjunction with decreased accessibility of melanocytic-phenotype genes including *MLANA* and *DCT*. Pathway analysis of the HDAC8-driven state showed an enrichment for pathways involved in focal adhesion, the cytoskeleton, proteoglycans, and multiple oncogenic signaling pathways (MAPK, PI3K, RAP1 and Ras). We reasoned that the NCSC state might be associated with increased metastatic potential and demonstrated that the HDAC8 expressing cells adopted an amoeboid-like phenotype that was associated with increased invasion into collagen and transendothelial cell migration. Injection of the isogenic HDAC8 expressing melanoma cells into the left ventricle of the heart increased the formation of brain metastases.

Metastasis of cancer cells to the brain is a complex, multi-step process [24, 26]. Although our studies did not recapitulate the entire brain metastatic cascade, we did model many key steps including 1) survival of the cells in the general circulation, 2) infiltration of the cells into the brain parenchyma and 3) the establishment of macrometastases. It is likely that the HDCA8-driven transcriptional state contributed to several of these processes. First, the effects of HDAC8 on the amoeboid cell state increased the robustness of the melanoma cells (e.g., the cells are more compact with increased mechanical strength) leading to their increased survival under fluid shear stress conditions. Second, the amoeboid phenotype increased invasive capacity allowing the cells to cross endothelial cell monolayers more efficiently and to migrate into thick collagen gels. A key feature of cancer cells that efficiently form brain metastases is increased expression of the Serpins, a family of protease inhibitors that help nullify the deleterious effects of the protease-rich brain microenvironment [25]. Analysis of our RNA-seq and ATAC-seq data supported a role for HDAC8 in driving the expression of multiple Serpins. In agreement with brain metastasis development being an end stage of melanoma progression [40], we found that increased expression of HDAC8 in clinical melanoma samples from the TCGA dataset was significantly associated with decreased overall survival. Interrogation of single cell RNA-seq data from a cohort of melanoma brain metastasis specimens revealed a correlation between expression of HDAC8 and the NCSC transcriptional signature.

HDACs exert their effects through deacetylation of both histone and non-histone targets. Our ChIP-seq and ATAC-Seq analyses demonstrated HDAC8 to have little effect upon global chromatin accessibility or promoter activity. Specific changes were however noted at discrete transcriptional sites, with in-depth analysis showing increased accessibility at AP-1, Jun, ATF3, FRA1 and TEAD sites. Less chromatin accessibility was seen at promoter sites for melanocytic-cell state associated transcription factors, such as MITF. ATAC-Seq analysis showed that accessibility to TEAD1 and TEAD4 binding sites increased upon expression of HDAC8 with ChIP-Seq demonstrating an initiation of transcription of TEAD-targeted genes. Many of the genes associated with the neural crest-like phenotype appear to be controlled by both TEAD and c-Jun transcription factors, indicating both transcription factor families may be under the regulation of HDAC8. These findings are supported by previous work showing that both AP-1/Jun and TEAD’s have also are important drivers of BRAFi resistance in melanoma [41, 42].

As ChIP-Seq analysis and immunoblotting for total histone acetylation did not suggest any global changes in histone acetylation following HDAC8 introduction we reasoned that non-histone targets of HDAC8 could be responsible for driving the NCSC transcriptional program we observed. We used mass spectrometry-based proteomics to map changes in protein acetylation and noted that introduction of HDAC8 altered the acetylation of multiple proteins including those involved in the cytoskeleton, metabolism, and DNA repair. Of relevance to the NCSC gene expression program, we also identified transcriptional co-regulators whose acetylation was modulated by HDAC8. One of the central hubs in the transcriptional regulator network was EP300 (E1A-binding protein, KAT3B) a histone acetyl transferase (HAT) with a high degree of homology to CREB-binding protein (CREBBP, CBP, KAT3A) [43, 44]. Both EP300 and CREBBP are HATs that primarily catalyze the acetylation of histone H3 at lysine 27 (H3K27ac) [44], which acts to mark super enhancer regions in the genome that form critical loci for the recruitment of master transcription factors. Increased expression of HDAC8 decreased H3K27 acetylation marks at the promoters of multiple melanocytic state genes including *MLANA* and *DCT*, and reduced chromatin access at melanocyte lineage transcription factor sites including MITF. While EP300 and CREBBP have similar homology and overlapping functions, they are also known to have distinct roles, depending upon the cellular context. In hematopoietic cells, CREBBP was required for proliferation and self-renewal, while EP300 was critical for differentiation and expression of genes important for cell cycle progression and DNA repair [45, 46]. H3K27ac ChIP-Seq studies have also shown distinct binding regions of both EP300 and CREBBP genome-wide, indicating EP300 and CREBBP regulate the enhancers of different genes [47, 48]. Our data suggest that HDAC8 selectively inhibits EP300 and not CREBBP, helping to explain why specific gene enhancer regions are affected without changes in global H3K27 acetylation.

In-depth proteomic analysis identified 3 lysine residues K1542, K1549, and K1550 in the HAT domain of EP300 that were deacetylated following introduction of HDAC8. EP300 HAT domain residues, including K1542 and K1549, have previously been shown to be deacetylated by the NAD^+^-dependent deacetylase sirtuin 2 (SIRT2), resulting in alterations in EP300 autoacetylation and protein binding [49]. Decreased acetylation of EP300 inhibited its HAT function in *in vitro* acetylation assays. Our data agree with previous studies showing that hypoacetylation of the EP300 activation loop, consisting of amino acids 1523-1554, decreases EP300 HAT activity [50].

Recent work has additionally suggested that EP300 directly acetylates MITF, leading to altered transcription factor distribution [31]. EP300 also directly acetylates c-Jun, leading to decreased DNA binding, increased turnover and decreased transcription of JUN targets [51, 52]. It is likely that the decrease in EP300 HAT activity following HDAC8 upregulation we observed would decrease MITF acetylation leading to a reduction of its transcriptional activity (as our ATAC-seq and RNA-seq data suggested). By contrast, inhibition of EP300 had opposite effects on Jun, with its deacetylation being associated with an increase in transcriptionally activated (phosphorylated) Jun and increased H3K27ac and chromatin accessibility at Jun promoters/genes. To further investigate the effects of EP300 inactivation in driving the NCSC phenotype we undertook phenotypic studies and found that both genetic and pharmacological inhibition of EP300 increased melanoma cell survival and mediated resistance of melanoma cells to BRAFi therapy. In addition to this, inhibition of EP300 also mimicked HDAC8 activation by promoting melanoma cell invasion and by increasing the expression of genes associated with the NCSC cell state.

We demonstrated for the first time that stress-mediated activation of HDAC8 leads to a switch from MITF to Jun driven transcriptional programs in melanoma cells that leads to the adoption of a NCSC state. The deacetylation of Jun increases its phosphorylation and transcriptional activity, leading to the initiation of an Jun/AP-1 driven, NCSC-like transcriptional program. At the same time, HDAC8-mediated deacetylation of EP300 inhibits its HAT function, decreasing accessibility at MITF promoter sites and altering the distribution of H3K27 at key melanocytic genes. Induction of this program by HDAC8 leads to the melanoma cells adopting a highly resilient, amoeboid phenotype that drives invasion and survival under both shear stress conditions and in the brain parenchyma. This HDAC8-driven program was also observed in melanocytes exposed to UV irradiation, providing the first link between a conserved melanocyte survival program and the aggressive metastatic behavior of melanoma cells.

## References

1. Tsao, H., et al., Melanoma: from mutations to medicine. Genes & Development, 2012. 26(11): p. 1131–1155.

2. Fife, K.M., et al., Determinants of outcome in melanoma patients with cerebral metastases. J Clin Oncol, 2004. 22(7): p. 1293–300.

3. Patchell, R.A., The management of brain metastases. Cancer Treatment Reviews, 2003. 29(6): p. 533–540.

4. Chen, G. and M.A. Davies, Emerging insights into the molecular biology of brain metastases. Biochemical Pharmacology, 2012. 83(3): p. 305–14.

5. Davies, M.A., et al., Integrated Molecular and Clinical Analysis of AKT Activation in Metastatic Melanoma. Clin Cancer Res, 2009. 15(24): p. 7538–7546.

6. Chen, G., et al., Molecular Profiling of Patient-Matched Brain and Extracranial Melanoma Metastases Implicates the PI3K Pathway as a Therapeutic Target. Clinical Cancer Research, 2014. 20(21): p. 5537–5546.

7. Bucheit, A.D., et al., Complete loss of PTEN protein expression correlates with shorter time to brain metastasis and survival in stage IIIB/C melanoma patients with BRAFV600 mutations. Clin Cancer Res, 2014. 20(21): p. 5527–36.

8. Cho, J.H., et al., AKT1 Activation Promotes Development of Melanoma Metastases. Cell Rep, 2015. 13(5): p. 898–905.

9. Brastianos, P.K., et al., Genomic Characterization of Brain Metastases Reveals Branched Evolution and Potential Therapeutic Targets. Cancer Discov, 2015.

10. Fischer, G.M., et al., Molecular Profiling Reveals Unique Immune and Metabolic Features of Melanoma Brain Metastases. Cancer Discov, 2019. 9(5): p. 628–645.

11. Tsoi, J., et al., Multi-stage Differentiation Defines Melanoma Subtypes with Differential Vulnerability to Drug-Induced Iron-Dependent Oxidative Stress. Cancer Cell, 2018. 33(5): p. 890–904 e5.

12. Rambow, F., et al., Toward Minimal Residual Disease-Directed Therapy in Melanoma. Cell, 2018.

13. Emmons, M.F., F. Faiao-Flores, and K.S. Smalley, The role of phenotypic plasticity in the escape of cancer cells from targeted therapy. Biochem Pharmacol, 2016.

14. Smalley, I., et al., Leveraging transcriptional dynamics to improve BRAF inhibitor responses in melanoma. EBioMedicine, 2019.

15. Konieczkowski, D.J., et al., A melanoma cell state distinction influences sensitivity to MAPK pathway inhibitors. Cancer Discov, 2014. 4(7): p. 816–27.

16. Emmons, M.F., et al., HDAC8 Regulates a Stress Response Pathway in Melanoma to Mediate Escape from BRAF Inhibitor Therapy. Cancer Res, 2019. 79(11): p. 2947–2961.

17. Lee, H., N. Rezai-Zadeh, and E. Seto, Negative regulation of histone deacetylase 8 activity by cyclic AMP-dependent protein kinase A. Mol Cell Biol, 2004. 24(2): p. 765–73.

18. Paraiso, K.H., et al., Ligand-Independent EPHA2 Signaling Drives the Adoption of a Targeted Therapy-Mediated Metastatic Melanoma Phenotype. Cancer Discov, 2015. 5(3): p. 264–73.

19. Schmidl, C., et al., ChIPmentation: fast, robust, low-input ChIP-seq for histones and transcription factors. Nat Methods, 2015. 12(10): p. 963–965.

20. Buenrostro, J.D., et al., Transposition of native chromatin for fast and sensitive epigenomic profiling of open chromatin, DNA-binding proteins and nucleosome position. Nat Methods, 2013. 10(12): p. 1213–8.

21. Smalley, I., et al., Single-Cell Characterization of the Immune Microenvironment of Melanoma Brain and Leptomeningeal Metastases. Clin Cancer Res, 2021. 27(14): p. 4109–4125.

22. Deardorff, M.A., et al., HDAC8 mutations in Cornelia de Lange syndrome affect the cohesin acetylation cycle. Nature, 2012. 489(7415): p. 313–7.

23. Hoek, K.S., et al., Metastatic potential of melanomas defined by specific gene expression profiles with no BRAF signature. Pigment Cell Res, 2006. 19(4): p. 290–302.

24. Phadke, M., et al., Melanoma brain metastases: Biological basis and novel therapeutic strategies. Exp Dermatol, 2022. 31(1): p. 31–42.

25. Valiente, M., et al., Serpins promote cancer cell survival and vascular co-option in brain metastasis. Cell, 2014. 156(5): p. 1002–16.

26. Kienast, Y., et al., Real-time imaging reveals the single steps of brain metastasis formation. Nat Med, 2010. 16(1): p. 116–22.

27. Fidler, I.J., Metastasis: quantitative analysis of distribution and fate of tumor emboli labeled with 125 I-5-iodo-2’-deoxyuridine. J Natl Cancer Inst, 1970. 45(4): p. 773–82.

28. Barnes, J.M., J.T. Nauseef, and M.D. Henry, Resistance to fluid shear stress is a conserved biophysical property of malignant cells. PLoS One, 2012. 7(12): p. e50973.

29. Taddei, M.L., et al., Mesenchymal to amoeboid transition is associated with stem-like features of melanoma cells. Cell Commun Signal, 2014. 12: p. 24.

30. Durbin, A.D., et al., EP300 Selectively Controls the Enhancer Landscape of MYCN-Amplified Neuroblastoma. Cancer Discov, 2022. 12(3): p. 730–751.

31. Louphrasitthiphol, P., et al., Tuning Transcription Factor Availability through Acetylation-Mediated Genomic Redistribution. Mol Cell, 2020. 79(3): p. 472–487 e10.

32. Haberland, M., et al., Epigenetic control of skull morphogenesis by histone deacetylase 8. Genes Dev, 2009. 23(14): p. 1625–30.

33. Haberland, M., R.L. Montgomery, and E.N. Olson, The many roles of histone deacetylases in development and physiology: implications for disease and therapy. Nat Rev Genet, 2009. 10(1): p. 32–42.

34. Long, J., et al., FLT3 inhibition upregulates HDAC8 via FOXO to inactivate p53 and promote maintenance of FLT3-ITD+ acute myeloid leukemia. Blood, 2020. 135(17): p. 1472–1483.

35. Dasgupta, T., et al., HDAC8 Inhibition Blocks SMC3 Deacetylation and Delays Cell Cycle Progression without Affecting Cohesin-dependent Transcription in MCF7 Cancer Cells. J Biol Chem, 2016. 291(24): p. 12761–12770.

36. Oehme, I., et al., Histone deacetylase 8 in neuroblastoma tumorigenesis. Clin Cancer Res, 2009. 15(1): p. 91–9.

37. Seto, E. and M. Yoshida, Erasers of histone acetylation: the histone deacetylase enzymes. Cold Spring Harb Perspect Biol, 2014. 6(4): p. a018713.

38. Yan, W., et al., Histone deacetylase inhibitors suppress mutant p53 transcription via histone deacetylase 8. Oncogene, 2013. 32(5): p. 599–609.

39. Hua, W.K., et al., HDAC8 regulates long-term hematopoietic stem-cell maintenance under stress by modulating p53 activity. Blood, 2017. 130(24): p. 2619–2630.

40. Davies, M.A., et al., Prognostic Factors for Survival in Melanoma Patients With Brain Metastases. Cancer, 2011. 117(8): p. 1687–1696.

41. Verfaillie, A., et al., Decoding the regulatory landscape of melanoma reveals TEADS as regulators of the invasive cell state. Nat Commun, 2015. 6: p. 6683.

42. Ramsdale, R., et al., The transcription cofactor c-JUN mediates phenotype switching and BRAF inhibitor resistance in melanoma. Sci Signal, 2015. 8(390): p. ra82.

43. Attar, N. and S.K. Kurdistani, Exploitation of EP300 and CREBBP Lysine Acetyltransferases by Cancer. Cold Spring Harb Perspect Med, 2017. 7(3).

44. Dhalluin, C., et al., Structure and ligand of a histone acetyltransferase bromodomain. Nature, 1999. 399(6735): p. 491–6.

45. Rebel, V.I., et al., Distinct roles for CREB-binding protein and p300 in hematopoietic stem cell self-renewal. Proc Natl Acad Sci U S A, 2002. 99(23): p. 14789–94.

46. Meyer, S.N., et al., Unique and Shared Epigenetic Programs of the CREBBP and EP300 Acetyltransferases in Germinal Center B Cells Reveal Targetable Dependencies in Lymphoma. Immunity, 2019. 51(3): p. 535–547 e9.

47. Ramos, Y.F., et al., Genome-wide assessment of differential roles for p300 and CBP in transcription regulation. Nucleic Acids Res, 2010. 38(16): p. 5396–408.

48. Martire, S., et al., Differential contribution of p300 and CBP to regulatory element acetylation in mESCs. BMC Mol Cell Biol, 2020. 21(1): p. 55.

49. Black, J.C., et al., The SIRT2 deacetylase regulates autoacetylation of p300. Mol Cell, 2008. 32(3): p. 449–55.

50. Thompson, P.R., et al., Regulation of the p300 HAT domain via a novel activation loop. Nat Struct Mol Biol, 2004. 11(4): p. 308–15.

51. Vries, R.G., et al., A specific lysine in c-Jun is required for transcriptional repression by E1A and is acetylated by p300. EMBO J, 2001. 20(21): p. 6095–103.

52. Zhang, D., T. Suganuma, and J.L. Workman, Acetylation regulates Jun protein turnover in Drosophila. Biochim Biophys Acta, 2013. 1829(11): p. 1218–24.

