## Supplemental Figures 1-13 for "HDAC8-mediated inhibition of EP300 drives a neural crest-like transcriptional state that increases melanoma brain metastasis"

### Slide 1
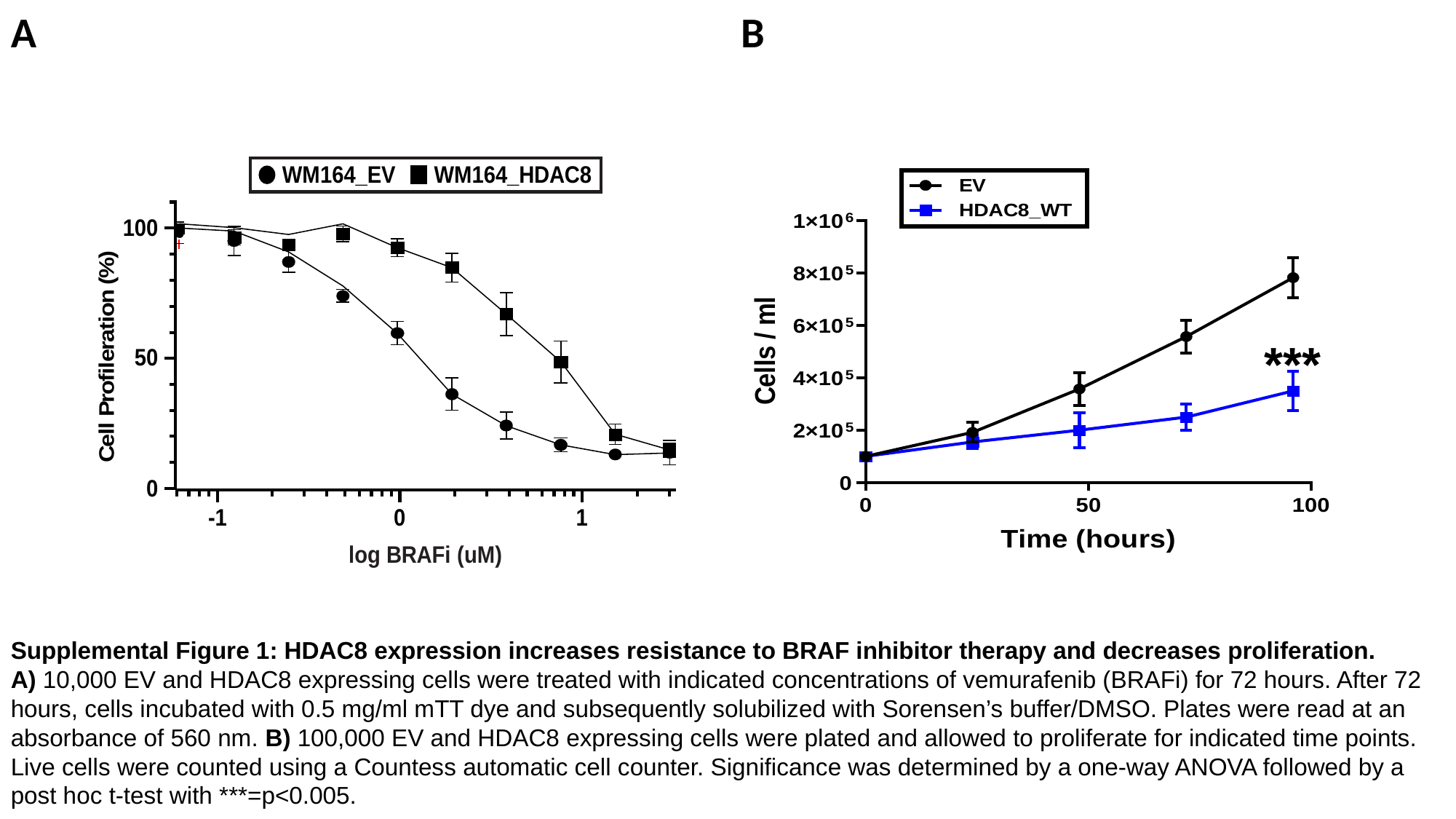

A
B
***
Supplemental Figure 1: HDAC8 expression increases resistance to BRAF inhibitor therapy and decreases proliferation.
A) 10,000 EV and HDAC8 expressing cells were treated with indicated concentrations of vemurafenib (BRAFi) for 72 hours. After 72 hours, cells incubated with 0.5 mg/ml mTT dye and subsequently solubilized with Sorensen’s buffer/DMSO. Plates were read at an absorbance of 560 nm. B) 100,000 EV and HDAC8 expressing cells were plated and allowed to proliferate for indicated time points. Live cells were counted using a Countess automatic cell counter. Significance was determined by a one-way ANOVA followed by a post hoc t-test with ***=p<0.005.

### Slide 2
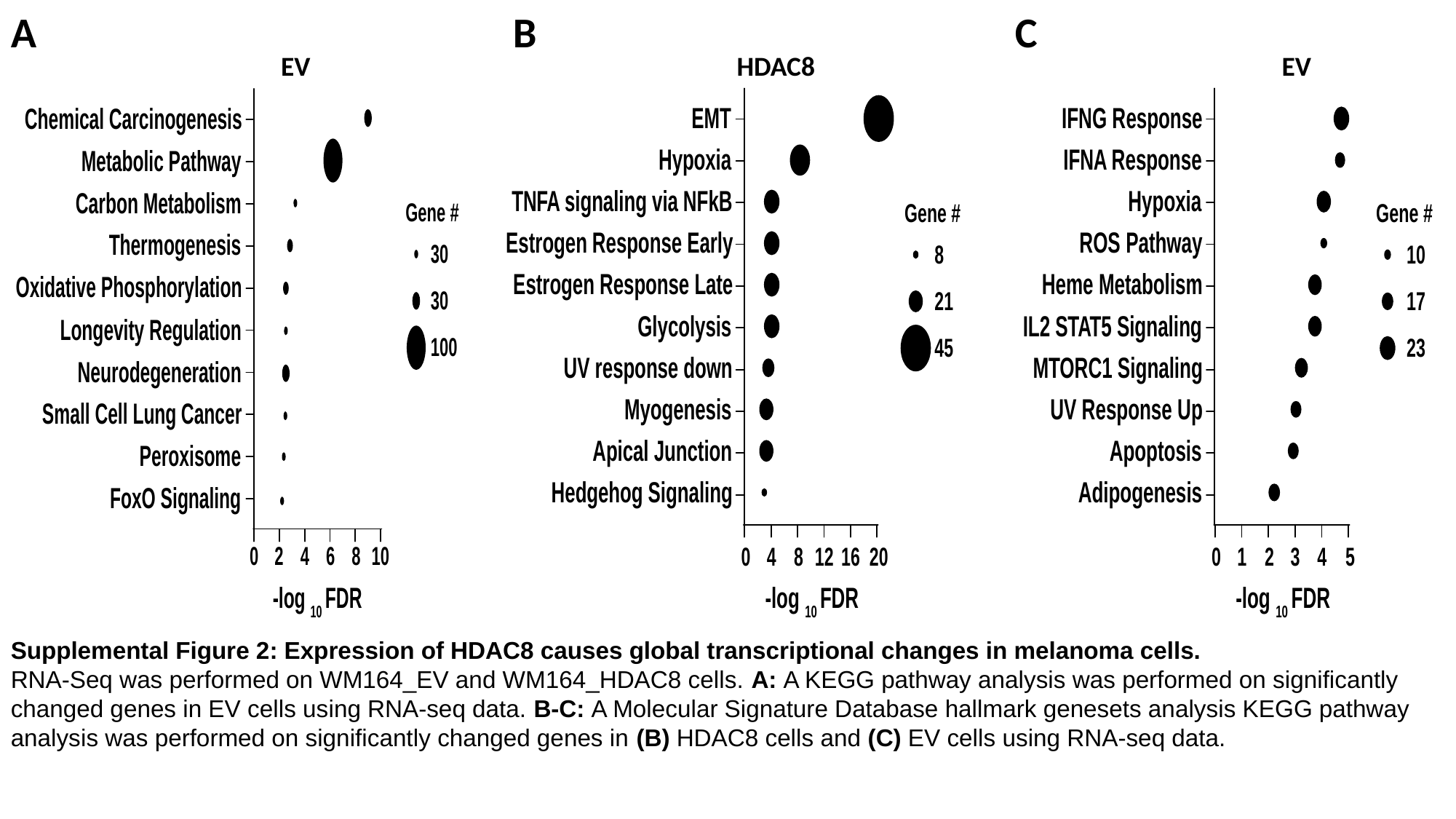

A
B
C
EV
HDAC8
EV
Supplemental Figure 2: Expression of HDAC8 causes global transcriptional changes in melanoma cells.
RNA-Seq was performed on WM164_EV and WM164_HDAC8 cells. A: A KEGG pathway analysis was performed on significantly changed genes in EV cells using RNA-seq data. B-C: A Molecular Signature Database hallmark genesets analysis KEGG pathway analysis was performed on significantly changed genes in (B) HDAC8 cells and (C) EV cells using RNA-seq data.

### Slide 3
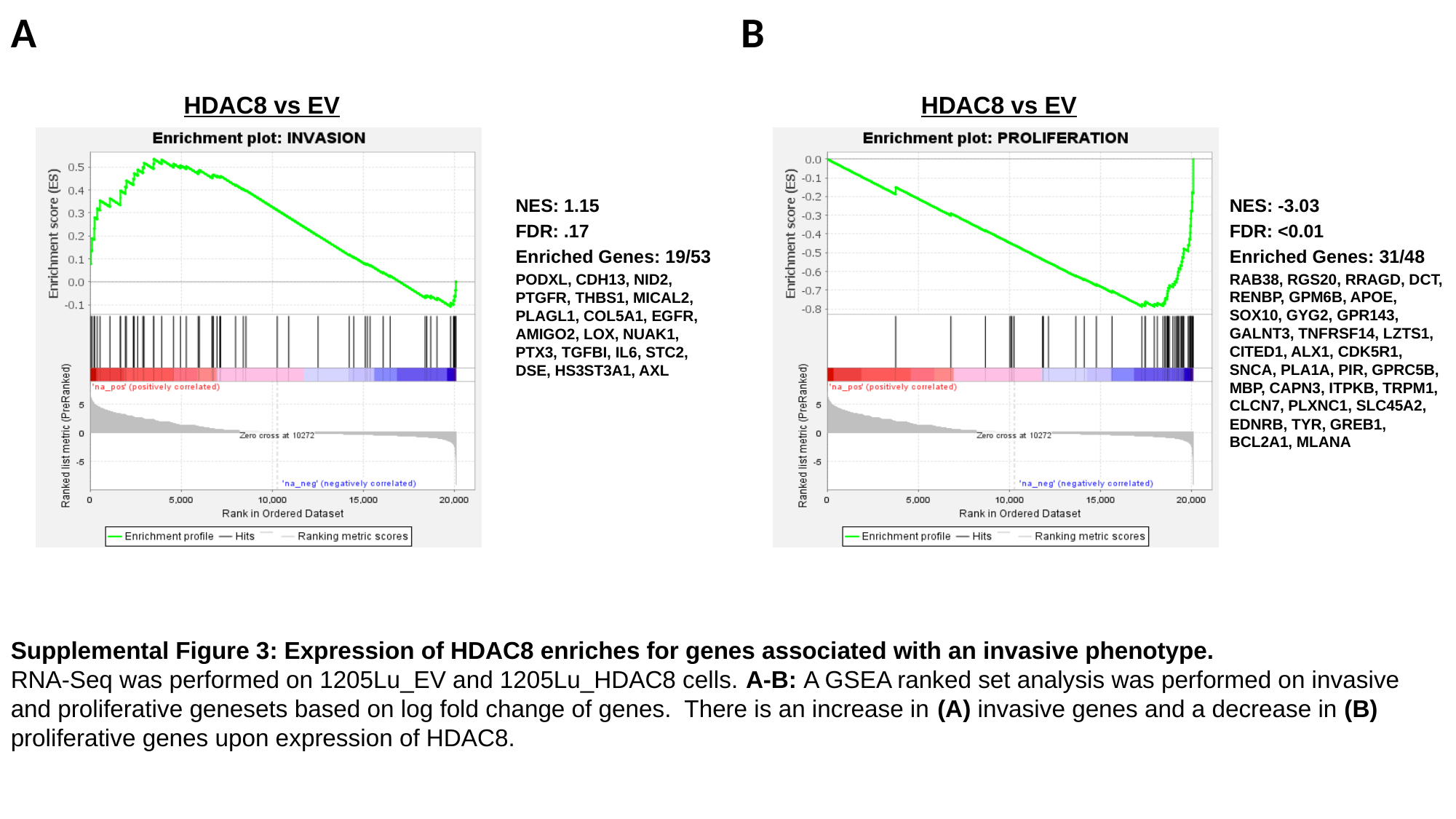

A
B
HDAC8 vs EV
HDAC8 vs EV
NES: 1.15
FDR: .17
Enriched Genes: 19/53
PODXL, CDH13, NID2, PTGFR, THBS1, MICAL2, PLAGL1, COL5A1, EGFR, AMIGO2, LOX, NUAK1, PTX3, TGFBI, IL6, STC2, DSE, HS3ST3A1, AXL
NES: -3.03
FDR: <0.01
Enriched Genes: 31/48
RAB38, RGS20, RRAGD, DCT, RENBP, GPM6B, APOE, SOX10, GYG2, GPR143, GALNT3, TNFRSF14, LZTS1, CITED1, ALX1, CDK5R1, SNCA, PLA1A, PIR, GPRC5B, MBP, CAPN3, ITPKB, TRPM1, CLCN7, PLXNC1, SLC45A2, EDNRB, TYR, GREB1, BCL2A1, MLANA
Supplemental Figure 3: Expression of HDAC8 enriches for genes associated with an invasive phenotype.
RNA-Seq was performed on 1205Lu_EV and 1205Lu_HDAC8 cells. A-B: A GSEA ranked set analysis was performed on invasive and proliferative genesets based on log fold change of genes. There is an increase in (A) invasive genes and a decrease in (B) proliferative genes upon expression of HDAC8.

### Slide 4
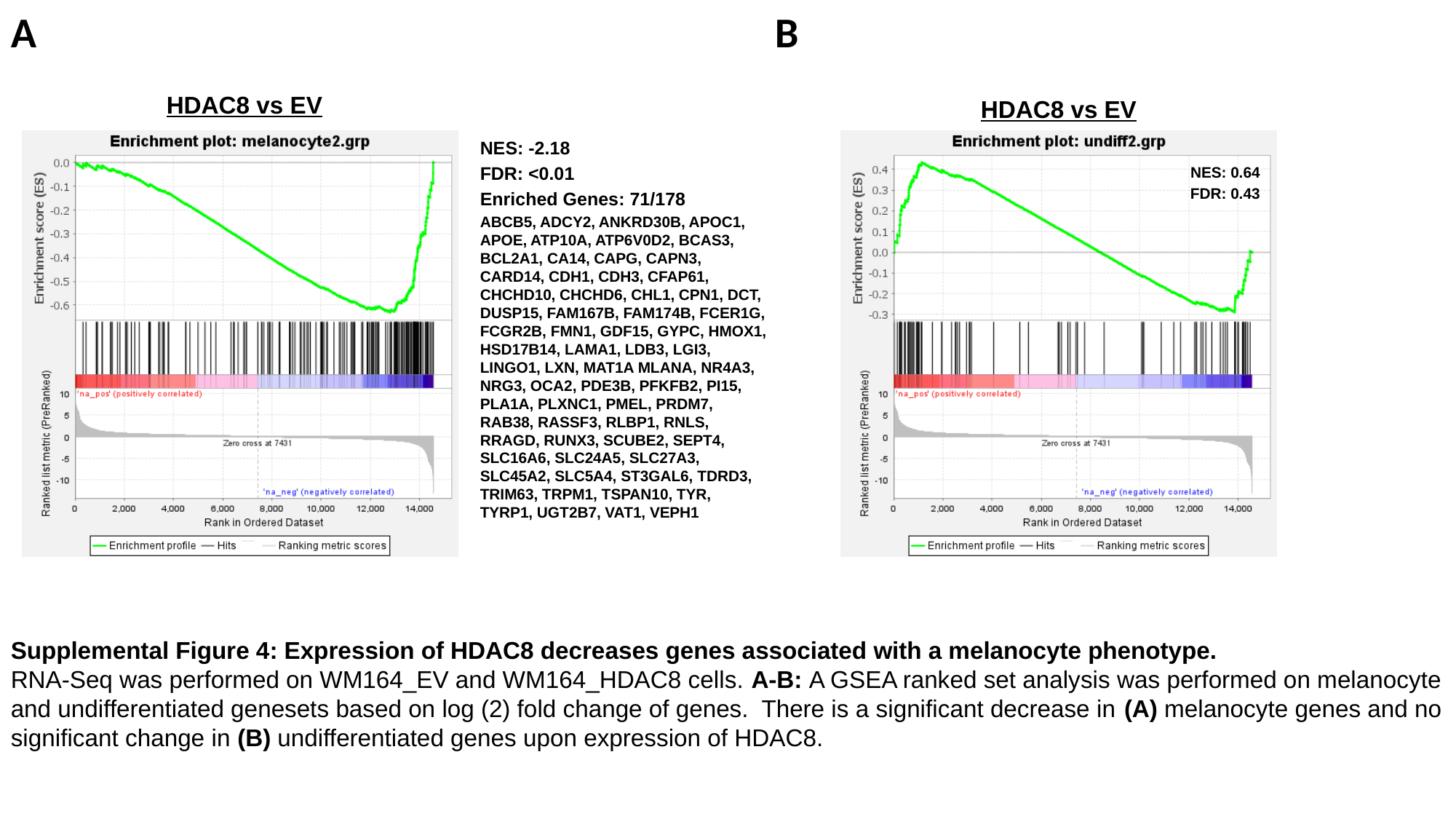

A
B
HDAC8 vs EV
HDAC8 vs EV
NES: -2.18
FDR: <0.01
Enriched Genes: 71/178
ABCB5, ADCY2, ANKRD30B, APOC1, APOE, ATP10A, ATP6V0D2, BCAS3, BCL2A1, CA14, CAPG, CAPN3, CARD14, CDH1, CDH3, CFAP61, CHCHD10, CHCHD6, CHL1, CPN1, DCT, DUSP15, FAM167B, FAM174B, FCER1G, FCGR2B, FMN1, GDF15, GYPC, HMOX1, HSD17B14, LAMA1, LDB3, LGI3, LINGO1, LXN, MAT1A MLANA, NR4A3, NRG3, OCA2, PDE3B, PFKFB2, PI15, PLA1A, PLXNC1, PMEL, PRDM7, RAB38, RASSF3, RLBP1, RNLS, RRAGD, RUNX3, SCUBE2, SEPT4, SLC16A6, SLC24A5, SLC27A3, SLC45A2, SLC5A4, ST3GAL6, TDRD3, TRIM63, TRPM1, TSPAN10, TYR, TYRP1, UGT2B7, VAT1, VEPH1
NES: 0.64
FDR: 0.43
Supplemental Figure 4: Expression of HDAC8 decreases genes associated with a melanocyte phenotype.
RNA-Seq was performed on WM164_EV and WM164_HDAC8 cells. A-B: A GSEA ranked set analysis was performed on melanocyte and undifferentiated genesets based on log (2) fold change of genes. There is a significant decrease in (A) melanocyte genes and no significant change in (B) undifferentiated genes upon expression of HDAC8.

### Slide 5
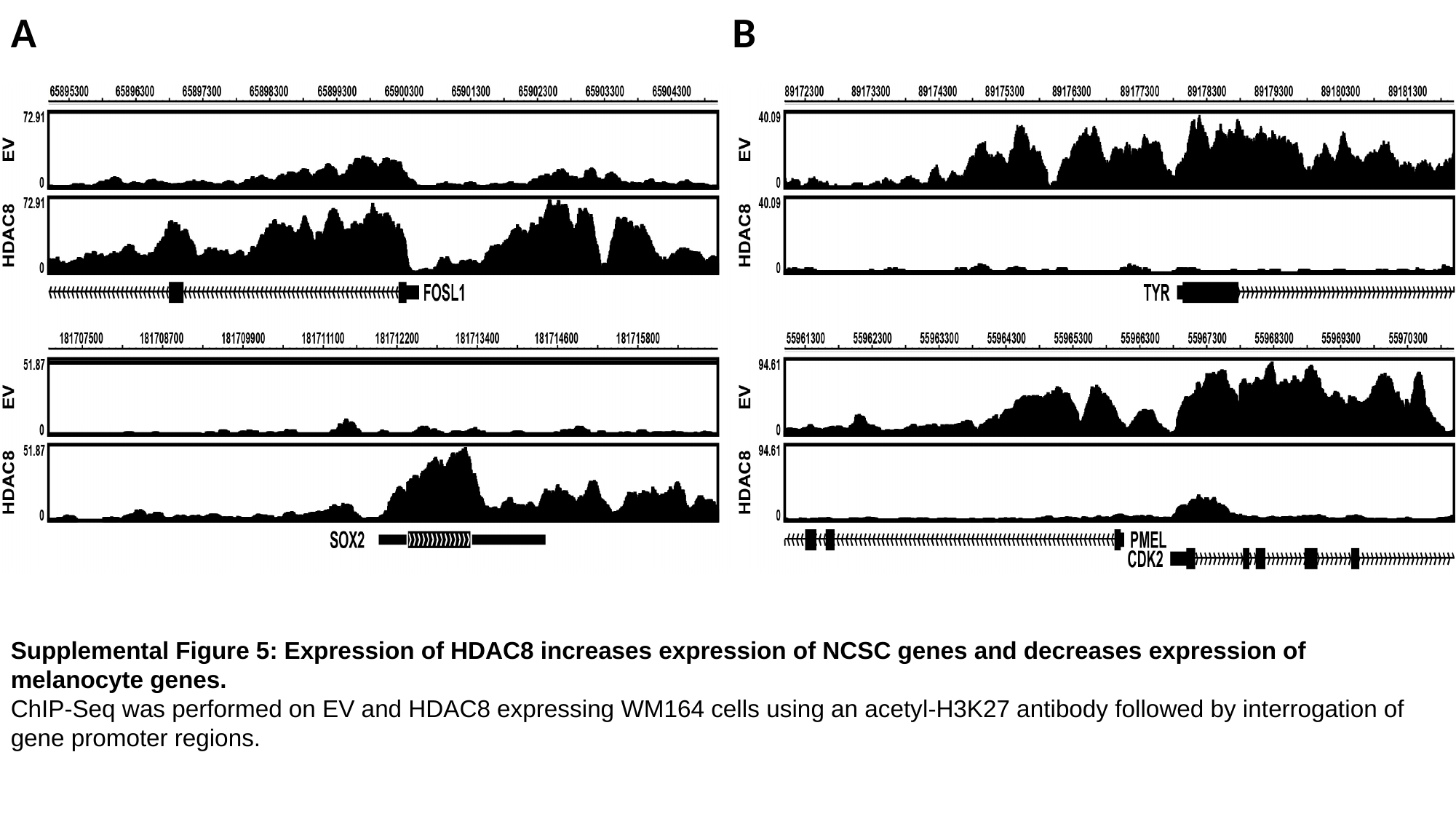

A
B
Supplemental Figure 5: Expression of HDAC8 increases expression of NCSC genes and decreases expression of melanocyte genes.
ChIP-Seq was performed on EV and HDAC8 expressing WM164 cells using an acetyl-H3K27 antibody followed by interrogation of gene promoter regions.

### Slide 6
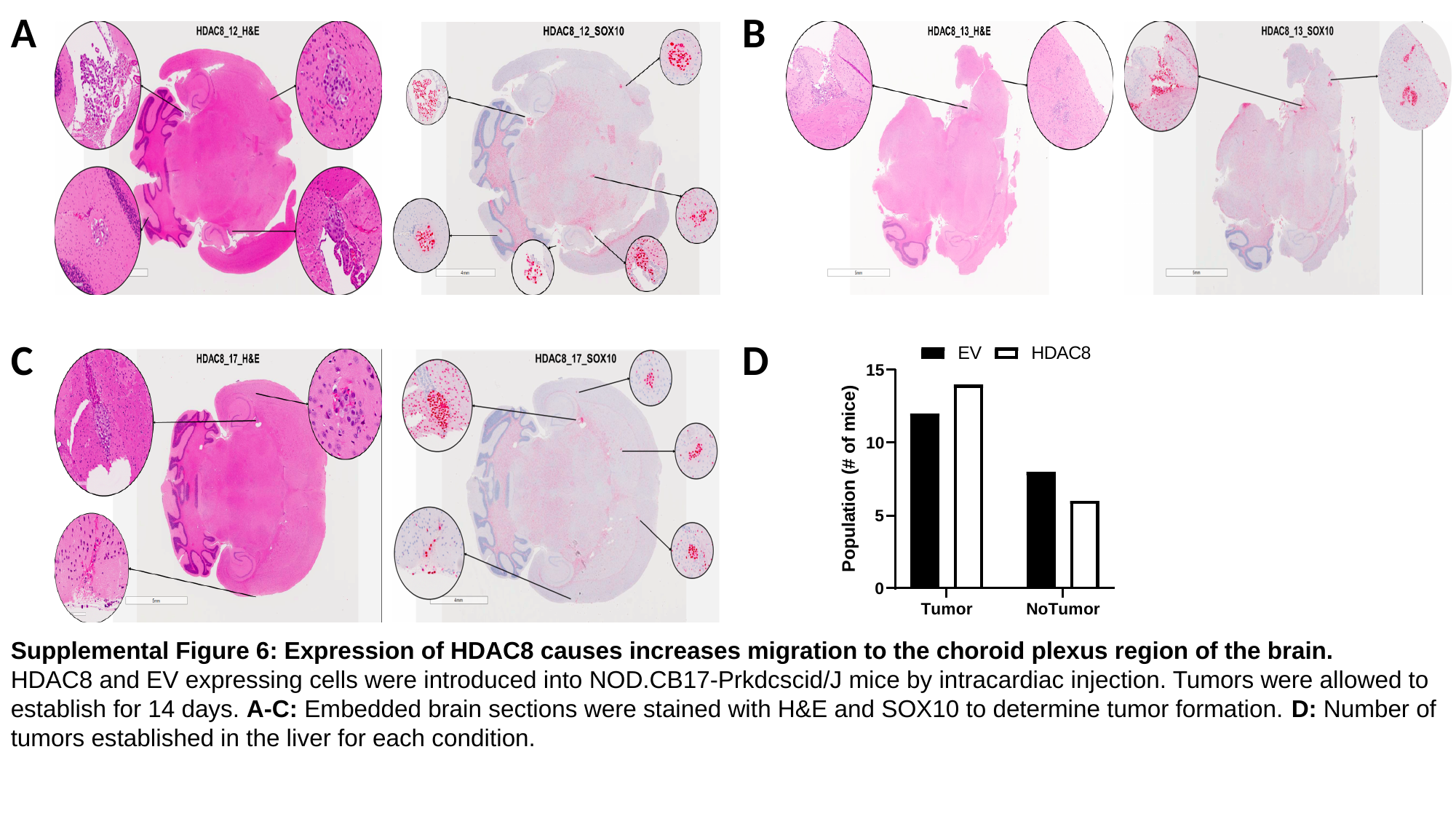

A
B
C
D
Supplemental Figure 6: Expression of HDAC8 causes increases migration to the choroid plexus region of the brain.
HDAC8 and EV expressing cells were introduced into NOD.CB17-Prkdcscid/J mice by intracardiac injection. Tumors were allowed to establish for 14 days. A-C: Embedded brain sections were stained with H&E and SOX10 to determine tumor formation. D: Number of tumors established in the liver for each condition.

### Slide 7
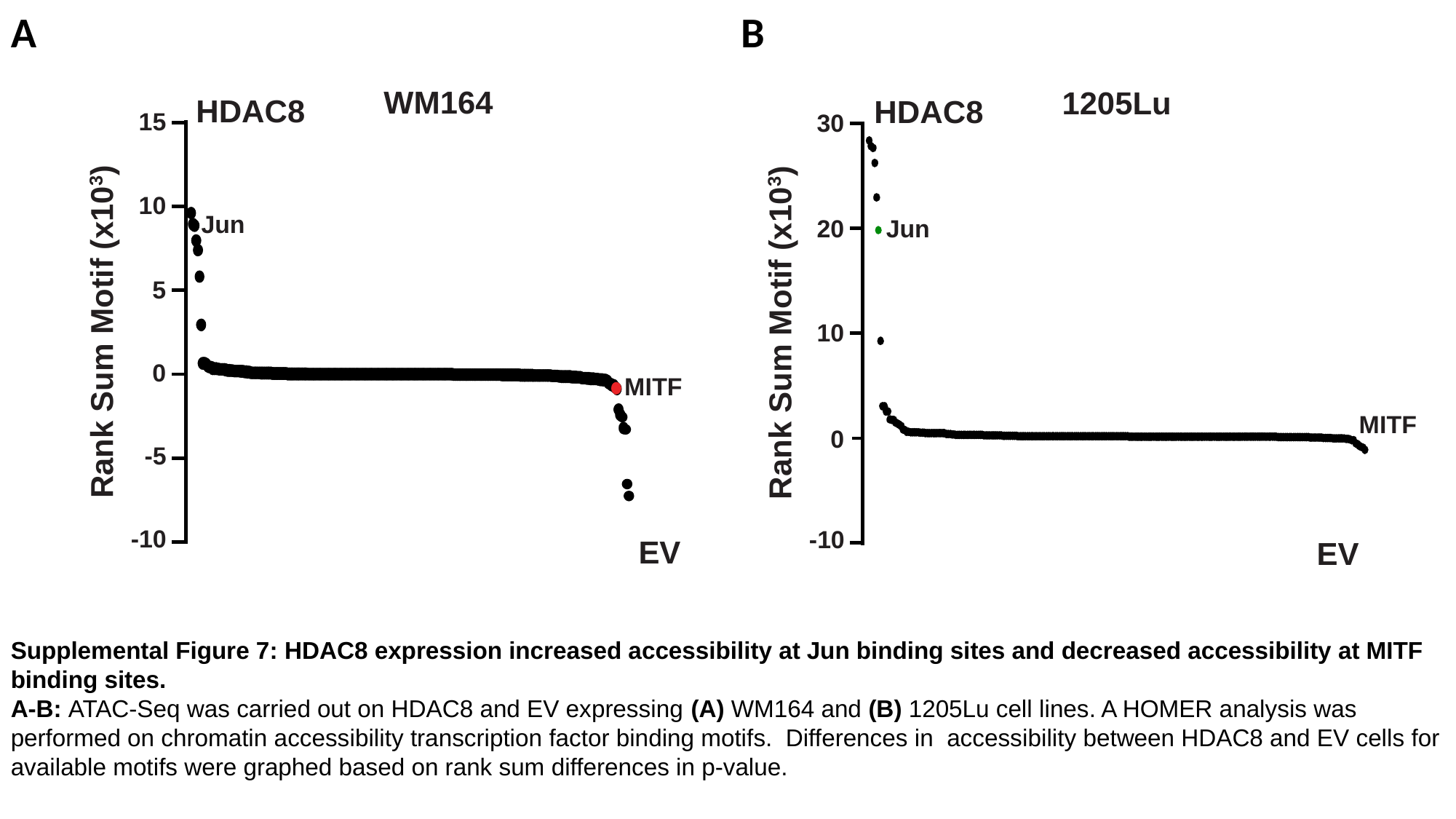

A
B
Supplemental Figure 7: HDAC8 expression increased accessibility at Jun binding sites and decreased accessibility at MITF binding sites.
A-B: ATAC-Seq was carried out on HDAC8 and EV expressing (A) WM164 and (B) 1205Lu cell lines. A HOMER analysis was performed on chromatin accessibility transcription factor binding motifs. Differences in accessibility between HDAC8 and EV cells for available motifs were graphed based on rank sum differences in p-value.

### Slide 8
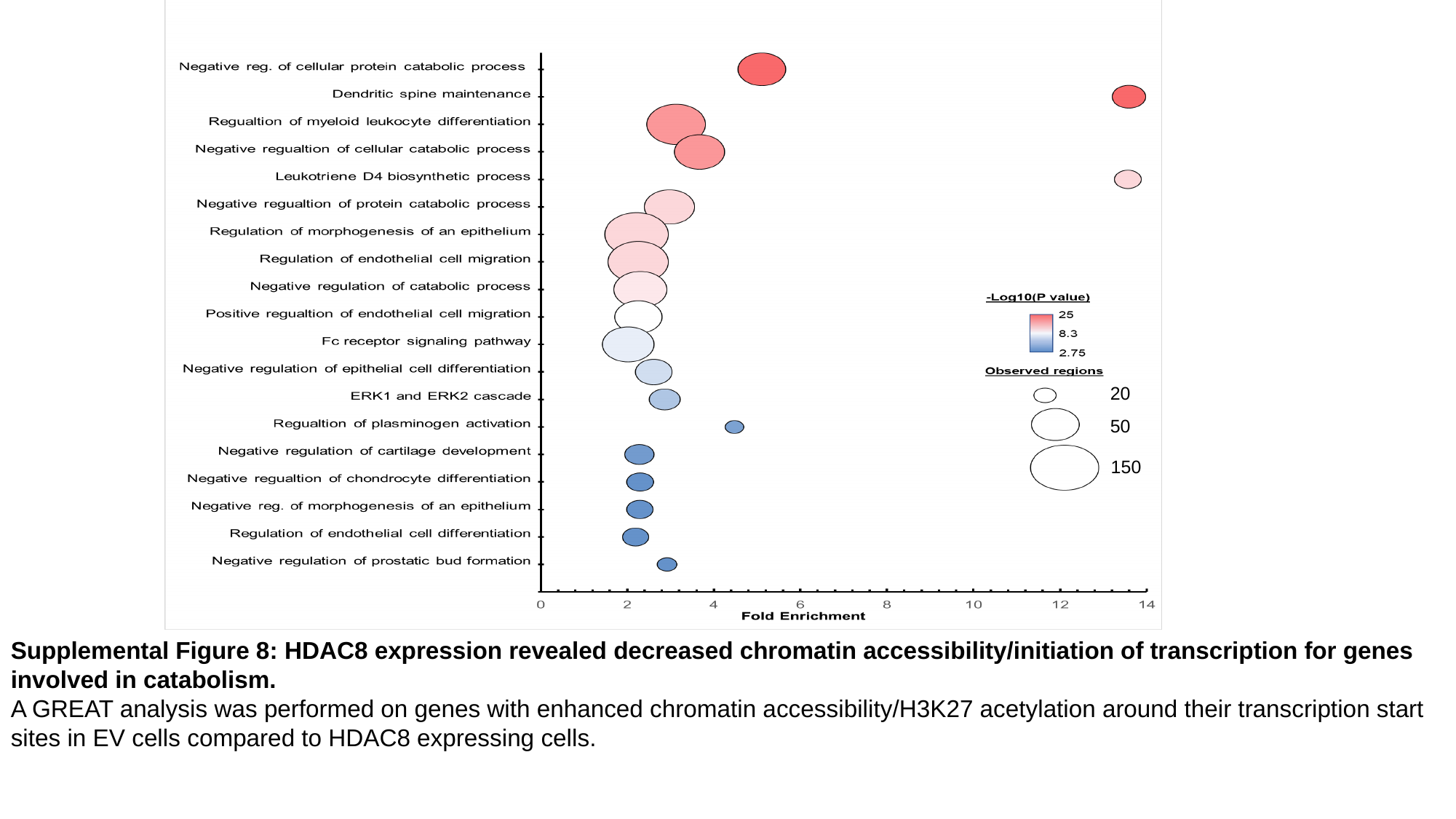

20
50
150
Supplemental Figure 8: HDAC8 expression revealed decreased chromatin accessibility/initiation of transcription for genes involved in catabolism.
A GREAT analysis was performed on genes with enhanced chromatin accessibility/H3K27 acetylation around their transcription start sites in EV cells compared to HDAC8 expressing cells.

### Slide 9
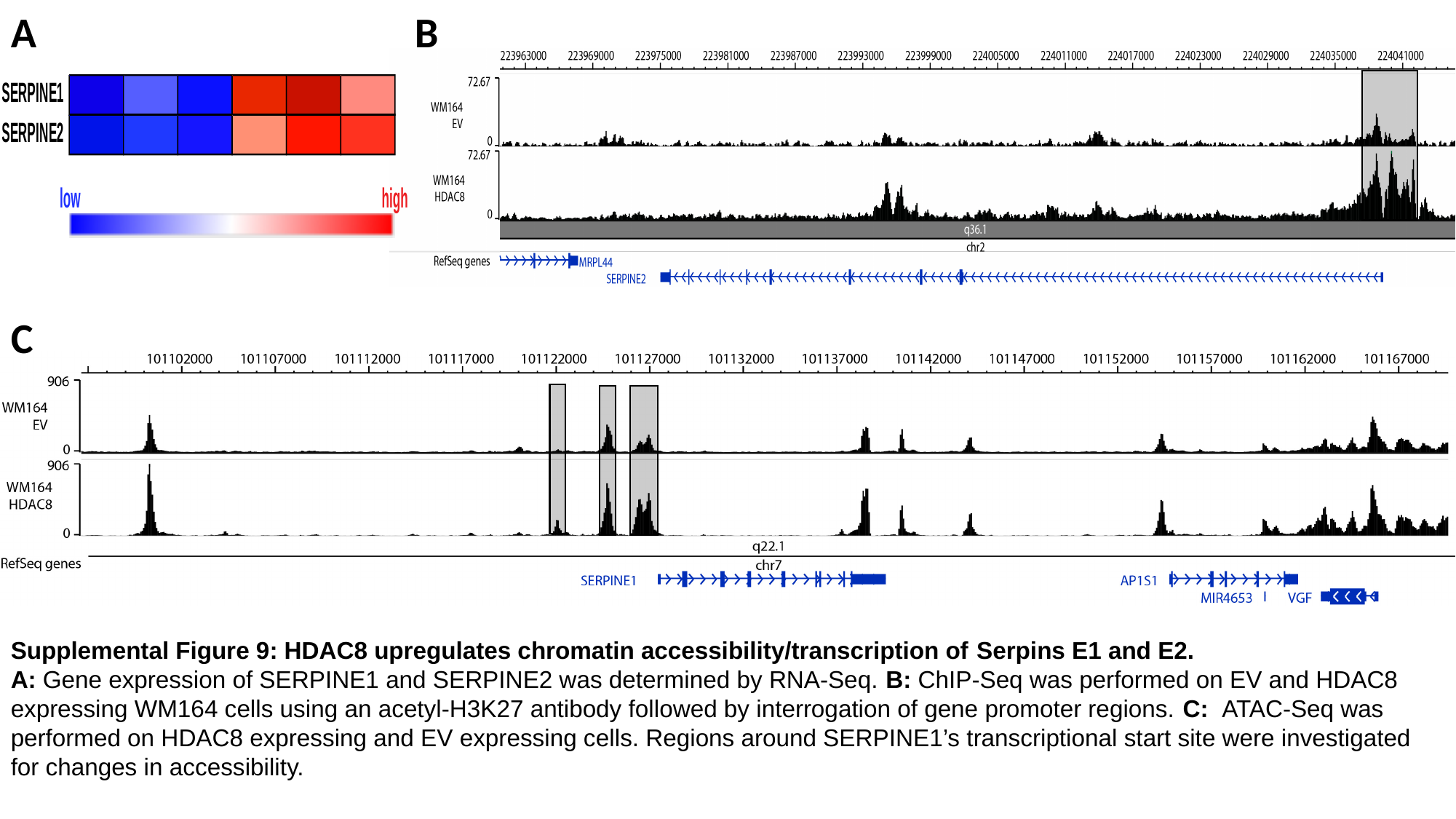

A
B
C
Supplemental Figure 9: HDAC8 upregulates chromatin accessibility/transcription of Serpins E1 and E2.
A: Gene expression of SERPINE1 and SERPINE2 was determined by RNA-Seq. B: ChIP-Seq was performed on EV and HDAC8 expressing WM164 cells using an acetyl-H3K27 antibody followed by interrogation of gene promoter regions. C: ATAC-Seq was performed on HDAC8 expressing and EV expressing cells. Regions around SERPINE1’s transcriptional start site were investigated for changes in accessibility.

### Slide 10
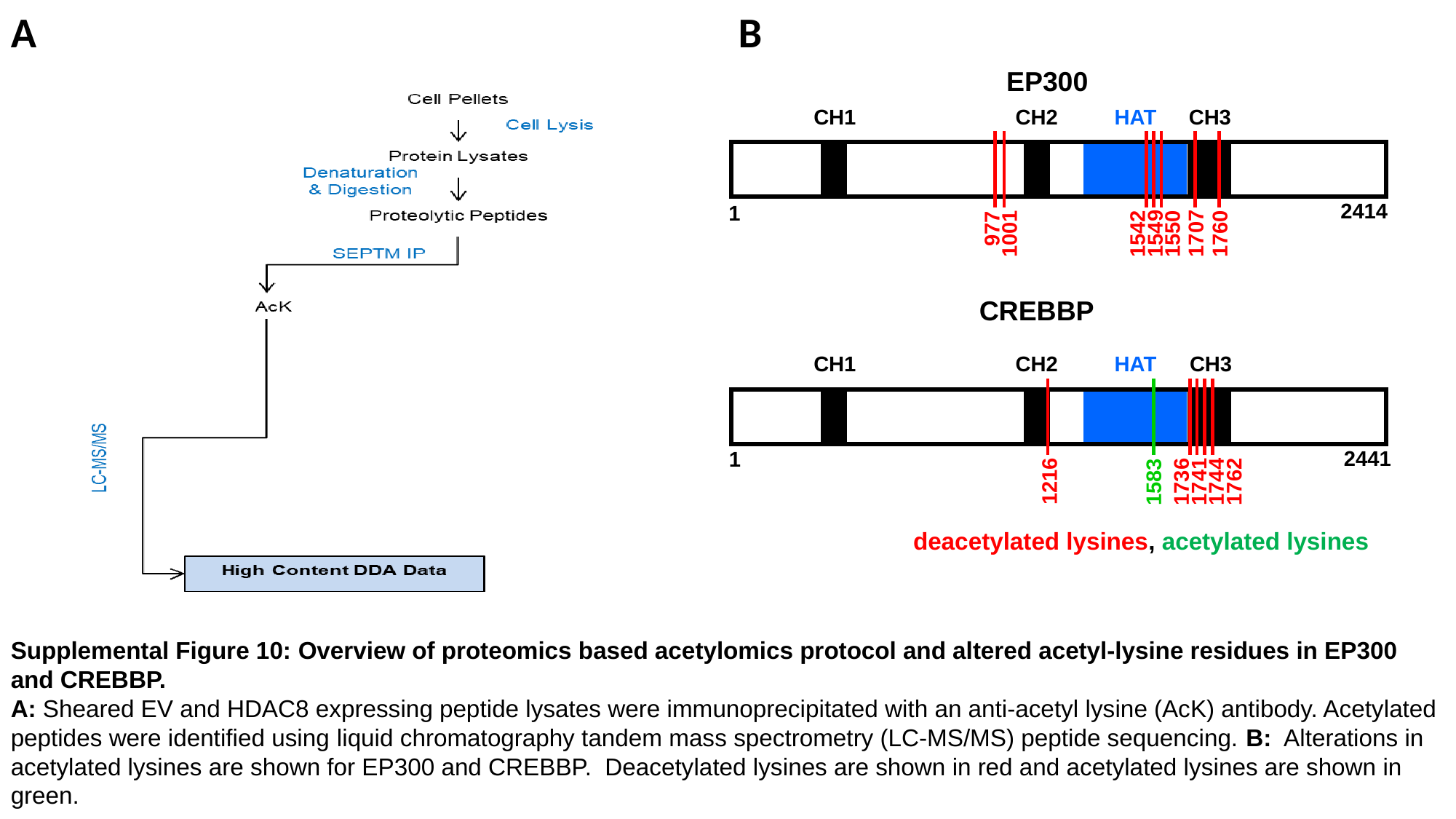

A
B
EP300
CH1
CH2
HAT
CH3
2414
1
977
1707
1549
1542
1550
1001
1760
CREBBP
CH1
CH2
HAT
CH3
2441
1
1216
1741
1762
1736
1744
1583
deacetylated lysines, acetylated lysines
Supplemental Figure 10: Overview of proteomics based acetylomics protocol and altered acetyl-lysine residues in EP300 and CREBBP.
A: Sheared EV and HDAC8 expressing peptide lysates were immunoprecipitated with an anti-acetyl lysine (AcK) antibody. Acetylated peptides were identified using liquid chromatography tandem mass spectrometry (LC-MS/MS) peptide sequencing. B: Alterations in acetylated lysines are shown for EP300 and CREBBP. Deacetylated lysines are shown in red and acetylated lysines are shown in green.

### Slide 11
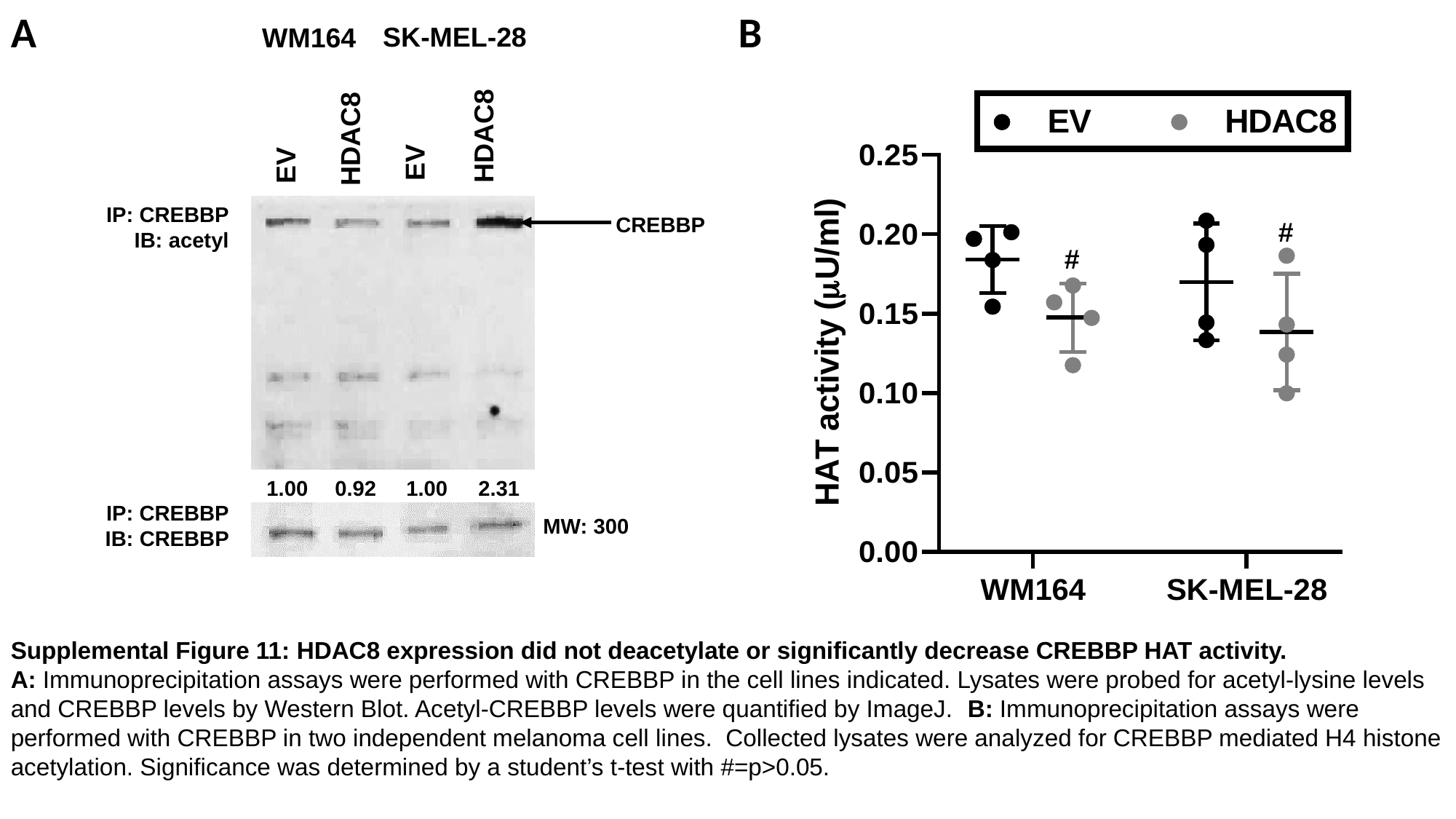

A
B
SK-MEL-28
WM164
EV
HDAC8
EV
HDAC8
IP: CREBBP
IB: acetyl
CREBBP
#
#
1.00
0.92
1.00
2.31
IP: CREBBP
IB: CREBBP
MW: 300
Supplemental Figure 11: HDAC8 expression did not deacetylate or significantly decrease CREBBP HAT activity.
A: Immunoprecipitation assays were performed with CREBBP in the cell lines indicated. Lysates were probed for acetyl-lysine levels and CREBBP levels by Western Blot. Acetyl-CREBBP levels were quantified by ImageJ. B: Immunoprecipitation assays were performed with CREBBP in two independent melanoma cell lines. Collected lysates were analyzed for CREBBP mediated H4 histone acetylation. Significance was determined by a student’s t-test with #=p>0.05.

### Slide 12
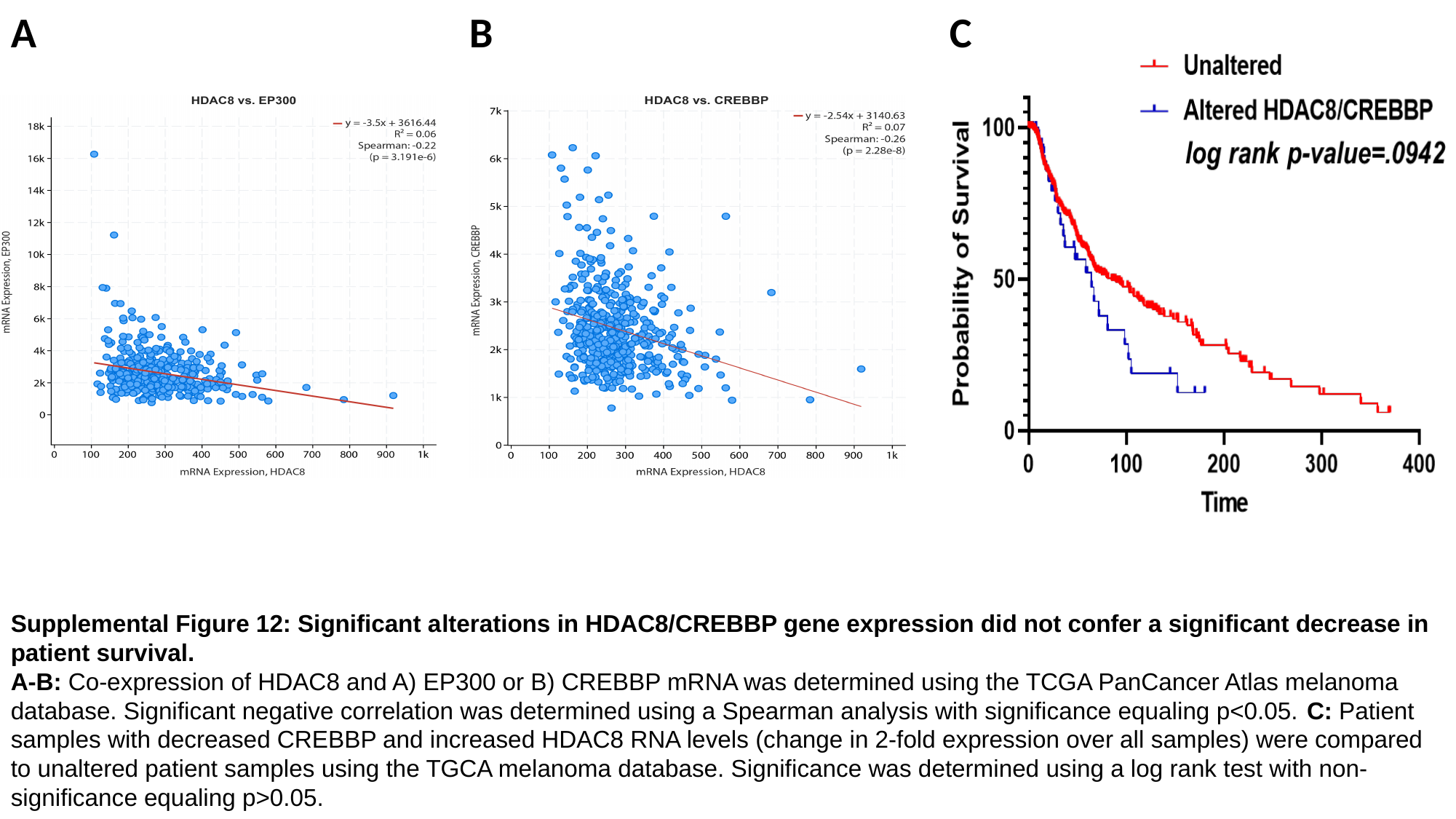

A
B
C
Supplemental Figure 12: Significant alterations in HDAC8/CREBBP gene expression did not confer a significant decrease in patient survival.
A-B: Co-expression of HDAC8 and A) EP300 or B) CREBBP mRNA was determined using the TCGA PanCancer Atlas melanoma database. Significant negative correlation was determined using a Spearman analysis with significance equaling p<0.05. C: Patient samples with decreased CREBBP and increased HDAC8 RNA levels (change in 2-fold expression over all samples) were compared to unaltered patient samples using the TGCA melanoma database. Significance was determined using a log rank test with non-significance equaling p>0.05.

### Slide 13
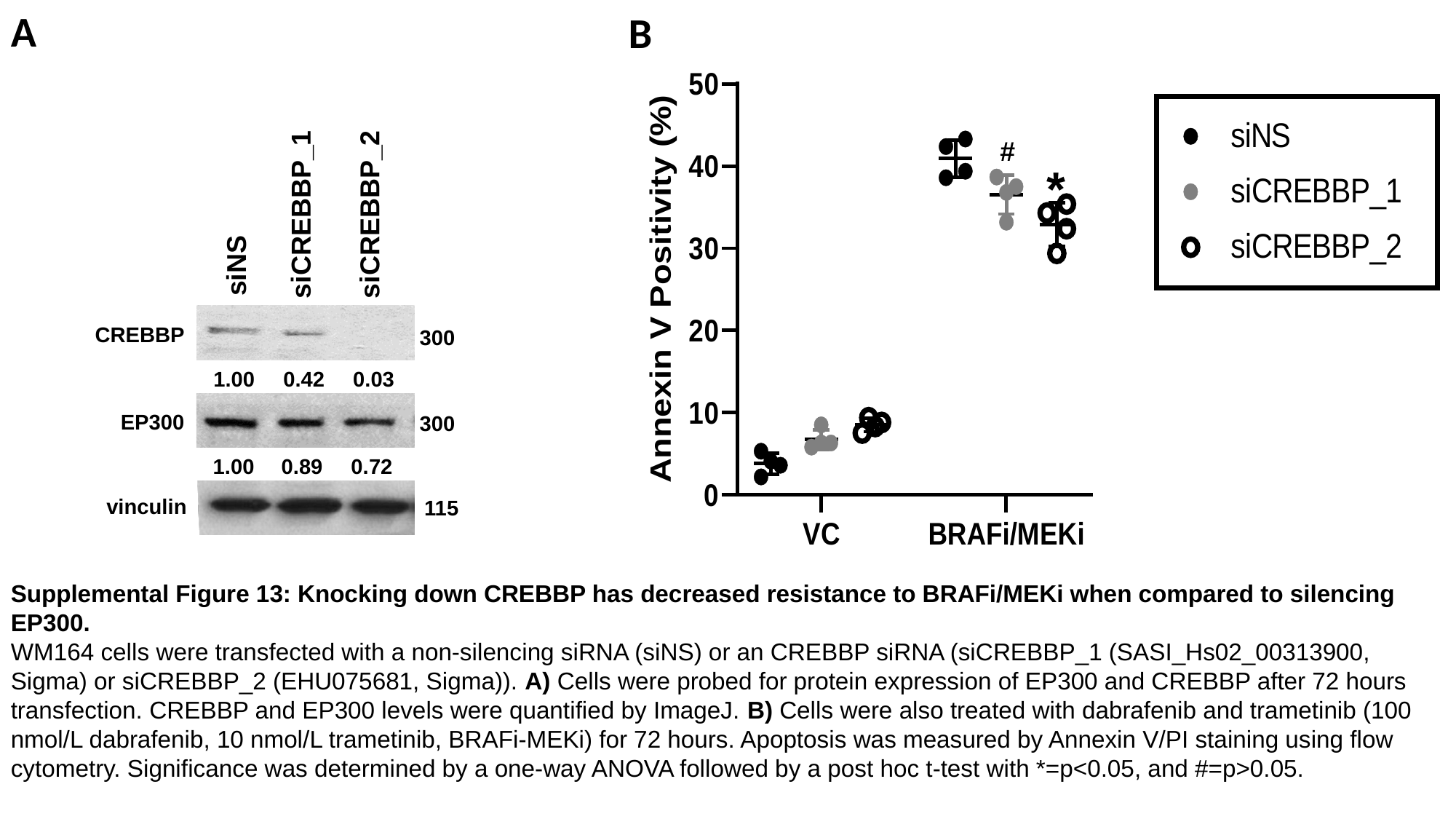

A
B
#
*
siCREBBP_1
siCREBBP_2
siNS
CREBBP
300
1.00
0.42
0.03
EP300
300
1.00
0.89
0.72
vinculin
115
Supplemental Figure 13: Knocking down CREBBP has decreased resistance to BRAFi/MEKi when compared to silencing EP300.
WM164 cells were transfected with a non-silencing siRNA (siNS) or an CREBBP siRNA (siCREBBP_1 (SASI_Hs02_00313900, Sigma) or siCREBBP_2 (EHU075681, Sigma)). A) Cells were probed for protein expression of EP300 and CREBBP after 72 hours transfection. CREBBP and EP300 levels were quantified by ImageJ. B) Cells were also treated with dabrafenib and trametinib (100 nmol/L dabrafenib, 10 nmol/L trametinib, BRAFi-MEKi) for 72 hours. Apoptosis was measured by Annexin V/PI staining using flow cytometry. Significance was determined by a one-way ANOVA followed by a post hoc t-test with *=p<0.05, and #=p>0.05.
